# Dilute phase oligomerization can oppose phase separation and modulate material properties of a ribonucleoprotein condensate

**DOI:** 10.1101/2021.04.19.440511

**Authors:** Ian Seim, Ammon E. Posey, Wilton T. Snead, Benjamin M. Stormo, Daphne Klotsa, Rohit V. Pappu, Amy S. Gladfelter

## Abstract

Ribonucleoprotein bodies are exemplars of membraneless biomolecular condensates that can form via spontaneous or driven phase transitions. The fungal protein Whi3 forms ribonucleoprotein condensates with different RNA molecules, and these condensates are implicated in key processes such as cell-cycle control and generating cell polarity. Whi3 has a modular architecture that includes a Q-rich intrinsically disordered region (IDR) and a tandem RNA recognition module. Here, we demonstrate that a 21-residue stretch within the Q-rich IDR has a weak intrinsic preference for forming alpha-helical conformations. Through mutagenesis, we find that increased alpha helicity enhances oligomerization in the dilute phase. One consequence of enhanced oligomerization is a dilution of Whi3 in the dense phase. The opposite behavior is observed when helicity within the 21-residue stretch of the Q-rich region is abrogated. Thus, the formation of dilute phase oligomers, driven by a specific sequence motif and potential synergies with the rest of the IDR, opposes incorporation of the Whi3 protein into the dense phase, thereby altering the dense phase stoichiometry of protein to RNA. Our findings, which stand in contrast to other systems where oligomerization has been shown to enhance the drive for phase separation, point to a novel mechanism that might be operative for influencing compositions of condensates. Our work also points to routes for designing synthetic ribonucleoprotein condensates whereby modulation of protein oligomerization via homotypic interactions can impact dense phase concentrations, stoichiometries, and material properties.

**Significance:** A large sub-class of biomolecular condensates are linked to RNA regulation and are known as ribonucleoprotein (RNP) bodies. While extensive work has identified driving forces for biomolecular condensate formation, relatively little is known about forces that oppose assembly. Here, using a fungal RNP protein, Whi3, we show that a portion of its intrinsically disordered, glutamine-rich region modulates phase separation by forming transient alpha helical structures that promote the assembly of dilute phase oligomers. These oligomers detour Whi3 proteins from condensates, thereby impacting the driving forces for phase separation, the protein-to-RNA ratio in condensates, and the material properties of condensates. Our findings show how nanoscale conformational and oligomerization equilibria can influence mesoscale phase equilibria.

## Introduction

Intracellular phase transitions are known to drive the formation and dissolution of membraneless biomolecular condensates (1). Prominent condensates include ribonucleoprotein (RNP) bodies such as stress granules (2–5) and processing bodies (5, 6) in the cytosol and nuclear bodies such as speckles (7) and nucleoli (8). RNP condensates can form via phase transitions that are influenced by a combination of homotypic and heterotypic protein-protein, protein-RNA, and RNA-RNA interactions (3, 4, 9–13).

Biomolecular condensates form through phase transitions known as phase separation aided percolation (PSAP) (14–17). The stickers-and-spacers framework provides a way to describe PSAP transitions (10, 14, 17–20). Phase separation is a density transition driven by non-specific, solvent-mediated interactions, whereas percolation is a networking transition driven by specific interactions among stickers (17, 21). Percolation transitions occur when protein and RNA molecules are topologically connected into system- or condensate-spanning networks. The solubility profiles of spacers contribute to density transitions, thereby determining whether percolation is aided by phase separation (17, 21).

Recent advances have identified the molecular features within protein (22–26) and RNA sequences (27, 28) that drive PSAP. These features include but are not restricted to: the number (valence) of folded stickers (16, 29, 30) such as RNA recognition motifs (RRMs) (20, 31) and protein-protein interaction domains (1); the valence and patterning of short linear motifs (stickers) within intrinsically disordered regions (IDRs) of proteins (7, 22, 24, 26, 32); the physical properties of spacers that are interspersed between stickers (14); and linear interaction motifs, sequencespecific secondary structures, the degree of unfoldedness, and purine versus pyrimidine contents within RNA molecules (4, 9, 10, 27, 28, 33).

Proteins that drive the formation of RNP condensates via PSAP often have modular architectures featuring structured oligomerization domains (SODs), IDRs, and RRMs (3). There are several examples of proteins that contain both IDRs and SODs. Prominent among these are the dimerization domain of the Nucleocapsid protein in SARS-CoV-2 (34, 35), the NTF2L dimerization domain of G3BP1/2 (2), a scaffold for stress granule formation, the oligomerization domain within coilin that drives Cajal Body formation (36), and the PB1 domain of auxin response factors (37) which gives rise to condensates that repress transcription during stress in plants. Oligomerization via SODs tethered to IDRs has also been shown to drive phase transitions in engineered systems (38–40). Synthetic and naturally occurring SODs are thought to act as sinks that lower the entropic penalty associated with phase transitions (39). In this view, SODs enable increased multivalence and the local concentrations of stickers that in turn drive phase transitions.

It is well known that many IDRs undergo coupled binding and folding reactions either through heterotypic interactions with cognate binding partners or via homotypic interactions that lead to dimers and higher-order oligomers (41–45). An archetypal example of dimerization induced folding of an IDR is the leucine zipper motif, which has a weak intrinsic preference for alpha-helical conformations and forms coiled-coil alpha-helical dimers (45, 46) or higher-order oligomers (47). Residues that define the interface among helices contribute to the oligomerization status of these leucine zipper systems. Overall, the stability of coiled coils is thermodynamically linked to the intrinsic preference for forming alpha-helices as monomers (48). How might sequence-specific preferences for alpha-helical conformations within IDRs and their oligomerization impact the phase behavior of RNA-binding proteins? Recent studies have identified the propensity to form alpha-helical structures within the C-terminal IDR of TDP-43 (49). This has been shown to be a key driver of phase separation (49). The findings for TDP-43 are reminiscent of findings regarding the effects of oligomerization driven by SODs. Accordingly, the consensus is that oligomerization via SODs enhances the driving forces for phase separation. Here, we uncover a surprising opposition to phase separation that derives from oligomerization driven by local ordering within an IDR.

The fungal protein Whi3 forms RNP condensates with different RNA molecules (9, 50). The simplest reconstituted system has two macromolecular components namely, the Whi3 protein and an RNA molecule such as the cyclin mRNA, *CLN3*. The Whi3 protein encompasses a C-terminal RRM and a long adjacent IDR that includes a glutamine-rich region (QRR) (**Fig. 1A**). Sequence analysis shows that the QRR contains a 21-residue motif, which we designate as CC. This CC motif has many of the features associated with alpha-helical coiled-coil forming domains (51). This opens the door to the possibility that the phase behavior of Whi3 is influenced by this putative helix- and coiled-coil-forming domain. Our work focuses on an interrogation of the effect of the CC motif on Whi3 phase behavior. We characterized the intrinsic helicity of the CC motif, in isolation, and assessed its impact on the phase behavior of full-length Whi3 through comparative studies of conformational and phase equilibria of mutants that diminish (CC^-^) or enhance (CC^+^) helicity. Surprisingly, we find that while enhancing intrinsic helicity enhances oligomerization in the dilute phase, it lowers the density of Whi3 in the RNP condensates. The converse is true when helicity is diminished. Spectroscopic investigations help establish the importance of disorder in the QRR for driving the intermolecular interactions that promote phase separation. Additionally, using coarse-grained simulations, we show how oligomerization and clustering within the dilute phase can modulate phase behavior in agreement with experimental observations. The main finding is that dilute phase oligomers act as off-pathway sinks for Whi3 proteins, whereas disorder within the QRR promotes phase separation via homotypic interactions.

**Figure 1:**
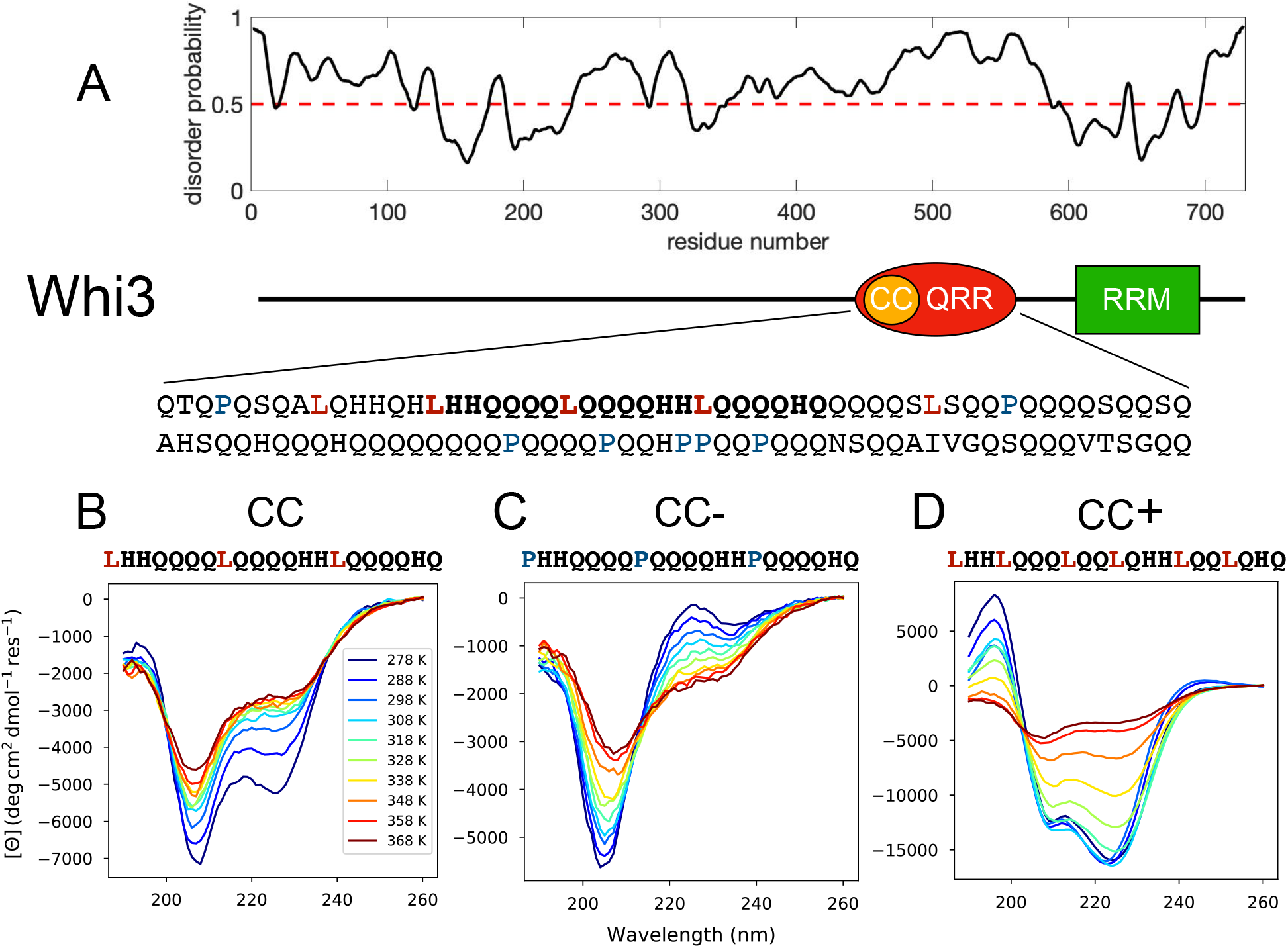
The QRR of Whi3 encompasses a putative CC motif. (A) Whi3 is predicted to be highly disordered (PrDOS) (60). The sequence includes a 108-residue QRR. (B-D) UV-CD spectra, measured at a series of temperatures, for the CC, CC^-^, and CC^+^ peptides, respectively. Peptide concentrations range from 600 – 900 μM.

## Results

### The QRR of Whi3 encompasses a region with a weak intrinsic preference for alphahelical conformations

The Whi3 protein encompasses a distinctive, 108-residue QRR that is predicted to be disordered (**Fig. 1A**). Sequence analysis revealed the presence of a 21-residue stretch, designated as CC, that has the characteristic features of domains that form alpha-helical coiled-coils (51, 52). These features include the absence of Pro and the presence of Leu at every seventh position (**Fig. 1A**). The putative CC motif appears to be an example of alpha-helical molecular recognition elements / features (a-MoREs or a-MoRFs) that are found in many IDRs (53, 54).

The sequence of the CC peptide is shown in **Fig. 1A**. We quantified the sequence-specificity of the alpha-helicity of the CC peptide by introducing point mutations that are likely to diminish / enhance helicity. The two designed peptides are designated as CC^-^ and CC^+^, respectively. The three Leu residues in the original CC peptide were replaced with Pro to generate CC^-^, a sequence variant designed to have diminished alpha helicity. To generate a variant designated as CC^+^, we replaced the fourth residue of each heptad repeat in the CC peptide with a Leu.

We performed ultraviolet circular dichroism (UV-CD) measurements for the three peptides (see methods for details). These spectra were collected at different temperatures at concentrations ranging from 600-900 μM; the data are shown in **Figs. 1B-D**. Across the temperature range of interest, the spectra for the CC peptide show two minima at 208 nm and 222 nm, respectively. The molar ellipticity is lower for the minimum at 208 nm. This feature is consistent with a heterogeneous ensemble of conformations albeit with alpha-helical character (55, 56). In contrast, and in accord with our predictions, each of the spectra for CC^-^ show only a single minimum at ~205 nm. There is a positive peak in the 215 – 218 nm range at low temperatures and this is concordant with the presence of Pro residues within the sequence (57). Overall, the spectra for CC^-^ are consistent with this peptide being disordered.

Finally, the spectra for the CC^+^ peptide demonstrate the characteristic minima at 208 nm and 222 nm. In addition, at temperatures below 338 K, the molar ellipticity at 222 nm is lower than the molar ellipticity at 208 nm. The molar ellipticities between 200 nm and 240 nm become less negative with increasing temperature and the presence of an isodichroic point at 200 nm is suggestive of CC^+^ showing a canonical helix-to-coil transition (58) as temperature increases. By fitting the molar ellipticity at 222 nm for each peptide to a two-state model, we estimated the melting temperatures for helix-to-coil transitions to be 333 K and 243 K for CC^+^ and CC, respectively (**Fig. S1**, *SI Appendix*). This implies that CC populates a heterogenous conformational ensemble at physiological temperatures, whereas CC^+^ is considerably more alpha-helical under these conditions. In contrast, CC^-^ has negligible preference for alpha-helical conformations.

We obtained an estimate of the secondary structure content of each peptide as a function of temperature using singular value deconvolution of the CD spectra via the CDSSTR algorithm, as implemented through the DichroWeb server (59). This analysis uses reference CD spectra of proteins with known structural content as basis sets from which to reconstruct the CD spectrum of interest. The accuracy of this approach is dependent on the extent to which the structural attributes of the protein of interest are represented in the basis set of proteins. Short peptides, coiled-coils, disordered regions, and aggregated or highly assembled states of proteins are either poorly represented or absent from these basis sets. Accordingly, we interpret the estimates of helical contents with a measure of caution. The fits are shown in **Fig. S2A-C**. The fractional helicities are highest at low temperatures and exhibit a single unfolding transition at high temperatures for CC and CC^+^. We performed forward and reverse temperature CD scans for the CC and CC^+^ peptides and did not find evidence of hysteresis in the fractional helicity for either peptide (**Fig. S2D, E**). Overall, the CD experiments suggest that the CC peptide has a partially helical character, CC^-^ has negligible helicity, and CC^+^ is strongly helical.

### The phase behavior of Whi3 is modulated by changes to the CC motif

To assess the impact of the CC motif on the phase behavior of Whi3, we studied the co-phase separation of Whi3 in the presence of the cyclin RNA *CLN3* (9) and compared this to the measured phase behavior obtained for three distinct variants. We expressed and purified full-length wild-type Whi3 (WT), and three variants namely, Whi3 (CC → CC^-^), Whi3 (CC → CC^+^), and Whi3 (ΔCC). In the Whi3 (ΔCC) protein, we deleted the 21 residues of the CC motif within Whi3. In each of our measurements, the Whi3 proteins and *CLN3* RNA molecules have fluorescent dye labels, and these are used to assess dense phase concentrations of protein and RNA molecules. In each mixture, 5% of the Whi3 is labeled and a single *CLN3* stock is used, such that the fluorescence values for both molecules can be directly compared across experiments.

All mutants of the Whi3 protein can still undergo phase separation with *CLN3*. However, the morphology and composition of assembled droplets varied amongst constructs. Notably, changes to the CC sequence resulted in significant differences in the dense phase stoichiometry (**Fig. 2A**). We quantified the concentrations of Whi3 in the dense phase by measuring fluorescence intensities and comparing to dye calibration curves (**Fig. S3A**, *SI Appendix*). When compared to condensates formed by WT Whi3 and *CLN3*, the condensates formed by Whi3 (ΔCC) have 1.5 times higher dense phase protein concentrations and Whi3 (CC → CC^-^) has 2 times the protein concentration in the dense phase. In contrast, the protein concentration in condensates formed by Whi3 (CC → CC^+^) is 0.2 times that of WT (**Fig. 2B**). This suggests that increased helicity in the CC motif leads to lower concentrations of Whi3 in the dense phase. Conversely, a disordered CC motif or deletion of the CC motif increases the dense phase concentration of Whi3.

**Figure 2:**
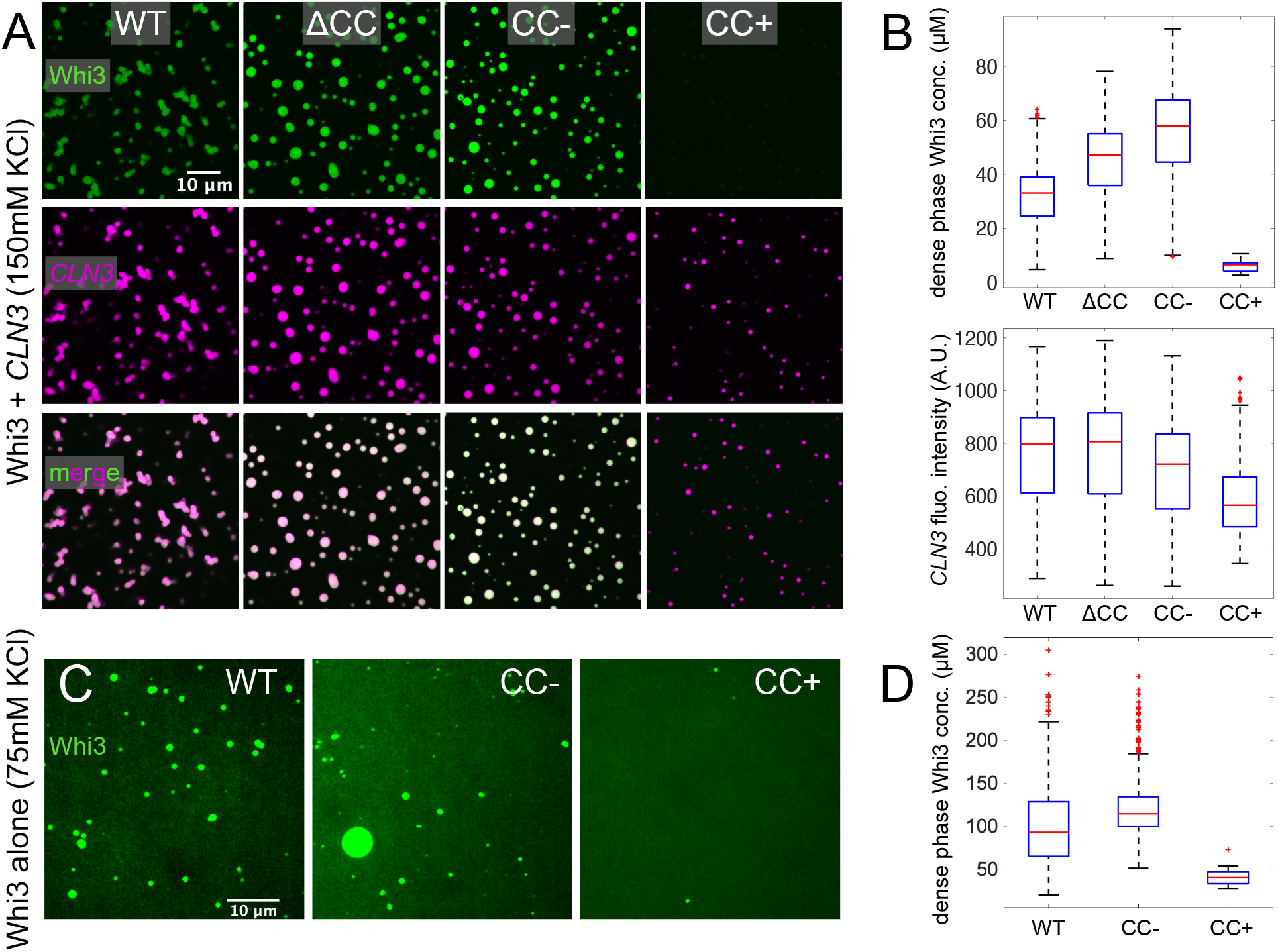
CC mutations alter *Whi3-CLN3* phase behavior. (A) Representative images of each protein at 5 hours after mixing 500 nM Whi3 with 5 nM *CLN3*. Whi3 (green) and *CLN3* (magenta) channels are contrasted identically in each image. (B) Box plots of dense phase Whi3 concentration and *CLN3* fluorescence intensity distributions of condensates in experiments corresponding to panel (A). Red lines indicate medians, edges of blue boxes indicate 25% and 75% quantiles, black lines at the edges of the dotted lines indicate the minimum and maximum observed values, and red stars indicate outliers. (C) Whi3-only phase separation at a bulk concentration of 17 μM recorded 4 hours after mixing. Images are contrasted such that fluorescence intensity is proportional to protein concentration. (D) Box plots of dense phase Whi3 concentration distributions for each condensate in experiments corresponding to panel (C).

An important consequence of the altered protein concentrations is a change in the ratio of protein to RNA that depends on the CC sequence. The dense phase concentrations of *CLN3* are similar for condensates formed by WT Whi3, Whi3 (ΔCC), and Whi3 (CC → CC^-^). However, the dense phase concentration of *CLN3* in condensates formed by co-phase separation of Whi3 (CC → CC^+^) with *CLN3* is about 0.8 times the dense phase concentration of *CLN3* in the WT system. Based on these results, we hypothesized that the effects of mutations within the CC motif should be evident even in the absence of *CLN3*. To test this hypothesis, we performed phase separation assays in the absence of *CLN3*. In line with results from the Whi3-*CLN3* system, we observe robust phase separation for Whi3 (CC → CC^-^), intermediate levels of phase separation for WT, and weak phase separation for Whi3 (CC → CC^+^) (**Fig. 2C, D**). In fact, at equivalent bulk protein concentrations of 17 μM, Whi3 (CC → CC^+^) forms very few droplets and those that form are very small (**Fig. 2D**). Relative to WT, the dense phase concentration increased by a factor of 1.2 for Whi3 (CC → CC^-^) whereas the Whi3 (CC → CC^+^) concentration was only 0.4 times that of WT. These results are qualitatively consistent with results from the Whi3-*CLN3* system. We conclude that homotypic interactions, driven by the CC motif, which we modulate using mutations, can alter the intrinsic phase behavior of Whi3 and its co-phase separation with *CLN3*.

### The CC motif enables sequestration of the Whi3 protein into dilute phase oligomers

The apparent opposition between oligomerization strength and the propensity to phase separate was unexpected. To arrive at a mechanistic understanding for these observations, we modeled the CC and QRR domains of Whi3 using the LASSI stickers and spacers model (18) (**Fig. 3A**). For simplicity, we focused on a minimal system comprising the CC-QRR fragment. We assume that CC is a dimerization motif that can access both dimerization competent and dimerization incompetent states. This assumption is motivated by the heterogeneous conformational distributions inferred from the CD measurements. In this model, the CC dimerizes with other CC domains via anisotropic interactions. This represents interactions between folded molecules. In addition, the CC motif functions like the rest of the QRR and engages in weak, isotropic interactions representing the disordered state. To represent the effects of mutations on the helicity and dimerization of the CC motif, we implement a trade-off between the energies associated with isotropic versus anisotropic interactions involving the beads that mimic the CC motif in the model (**Fig. 3A**). The right-most system in **Fig. 3** represents the extreme case that is most like Whi3 (CC → CC^+^), with stable helices enabling strong anisotropic binding. The central system corresponds to the opposite limit, mimicking Whi3 (CC → CC^-^), with a completely unfolded CC motif that interacts purely via isotropic interactions. The WT system has a CC bead that interacts via a combination of weaker anisotropic and isotropic interaction energies.

**Figure 3:**
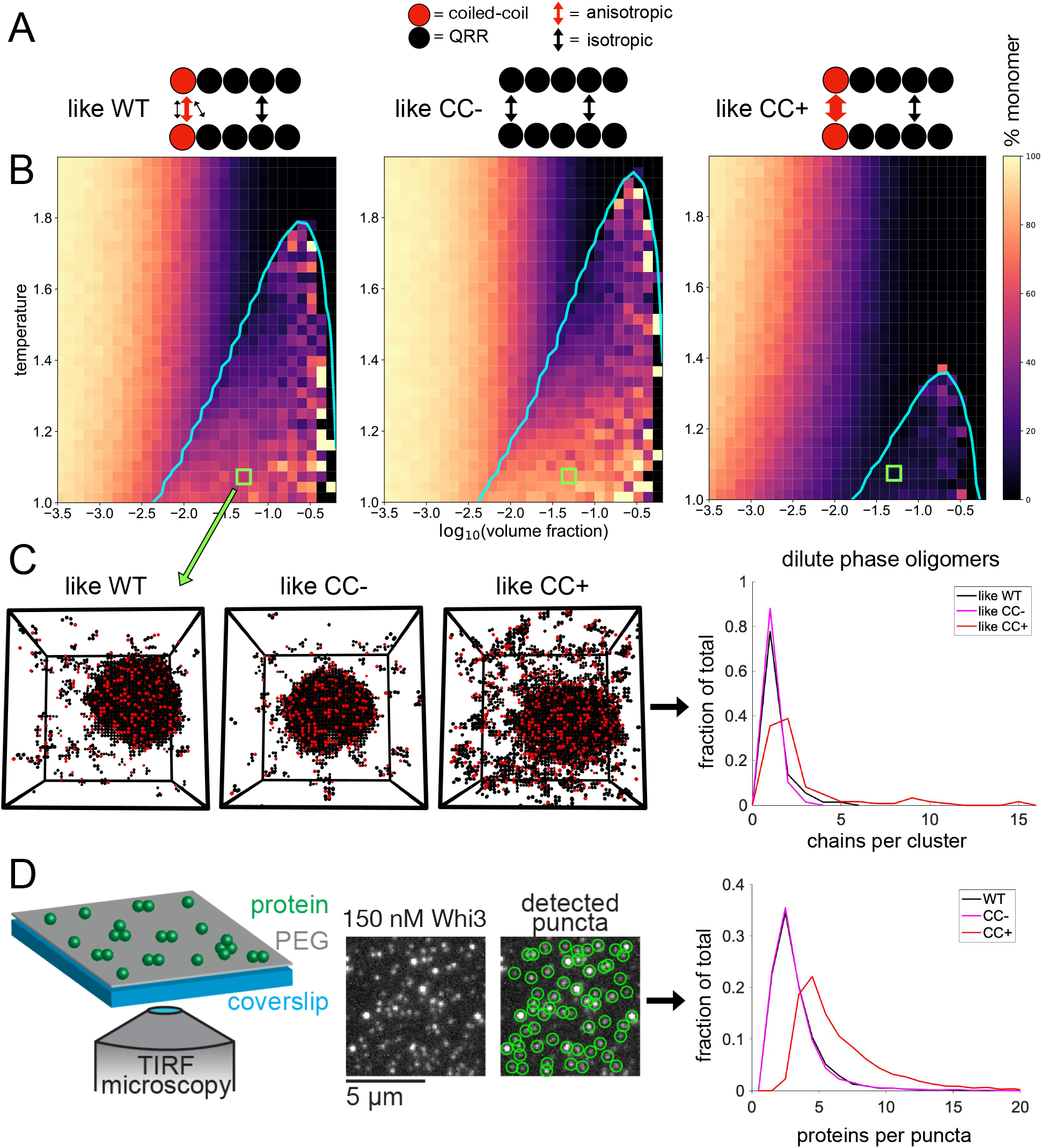
Oligomerization weakens phase separation and vice versa. (A) The CC-QRR fragment is modeled as five beads, with each bead corresponding to ~20 residues. The thickness of arrows corresponds to the strength of interactions. (B) Plots show binodals (cyan) overlaid on heatmaps indicating the percentage of the dilute phase / homogeneous phase that is monomeric. (C) Snapshots of simulations from regions corresponding to green squares in (B) are shown, and the dilute phase oligomer distributions are quantified for these snapshots. (D) Whi3 dilute phase cluster sizes at low concentrations are quantified via TIRF microscopy and particle detection.

For each system, we performed a series of LASSI simulations across a range of concentrations and temperatures. We computed coexistence curves using the approach prescribed by Choi et al. (18). The computed phase boundaries show that changes to the interactions mediated by the CC bead are sufficient to alter the driving forces for phase separation (**Fig. 3B**). We observe a suppression of phase separation when the isotropic interactions are weakened in favor of strengthening the anisotropic interactions. The converse is true when the isotropic interactions are strengthened over anisotropic interactions. This behavior is consistent with results from measurements that were performed in the absence of *CLN3* (**Fig. 2D**).

For each of the simulations, we quantified the distributions of cluster sizes in the coexisting dilute phases when phase separation occurred, and for the homogeneous system that lies outside the phase boundary. Enhancement of CC dimerization contributes to an increase in the sizes of dilute phase oligomers inside and outside the two-phase regime (**Fig. 3C**). Although the CC beads can only form dimers, they are connected to QRRs within the chain molecules. Accordingly, increasing the strength of anisotropic interactions mediated by the CC increases the local concentration of appended QRRs, leading to enhanced interactions among QRRs. This leads to large, dilute phase oligomers that act as sinks for Whi3 molecules. These sinks weaken the driving forces for Whi3 phase separation and may be viewed as “off-pathway” species as far as Whi3 phase separation is concerned.

To test predictions from the LASSI simulations, we measured oligomer sizes for WT, Whi3 (CC → CC^-^), and Whi3 (CC → CC^+^) at low concentrations that correspond to the one-phase regime (**Fig. 3D**). Using the fluorescence of single Whi3 molecules as a calibration, we recorded the distribution of oligomer sizes for each protein. Our predictions from LASSI-based simulations are supported by the observation that, despite their greatly reduced phase separation ability, Whi3 (CC → CC^+^) proteins form a heterogeneous distribution of oligomers in the dilute phase.

Taken together, our results indicate that the helicity-enabled oligomerization, driven by the CC motif, can help sequester the Whi3 protein into the dilute phase, thereby impacting the driving forces for phase separation and altering the dense phase Whi3 concentration. By changing the helicity of CC, the balance of protein between the two phases is shifted. This is observed in the different Whi3-*CLN3* systems, indicating that the protein density in Whi3-RNA condensates is impacted, in part, by homotypic associations among Whi3 molecules that give rise to finite-sized clusters in the dilute phase.

### Evidence for loss of helicity driving phase separation

Our data suggest that helix formation within the CC motif is linked to oligomerization that enables sequestration of Whi3 molecules in dilute phase oligomers that form via CC-mediated interactions and are further stabilized by interactions among QRRs. In contrast, a lack of helicity within the CC motif drives phase separation, which leads to higher protein concentrations in dense phases characterized by non-stoichiometric assemblies. Inspired by the work of Urry and coworkers (61, 62) and that of Duysens (63), we tested the hypothesis that loss of helicity through the CC motif drives phase separation. For this, we turned to UV-CD measurements and collected spectra as a function of peptide / protein concentration at room temperature. The constructs studied include the three peptides CC, CC^-^, and CC^+^ as well as the full-length Whi3 WT protein. Systems that undergo phase separation are known to show a strong distortion of the CD spectra that consists of a flattening of the spectra and the appearance of a new minimum at ~230 nm (63). Urry and coworkers (61, 62) noted that these features of CD spectra are related to the increased sizes of particles in solution with increasing protein concentration. Accordingly, we used UV-CD spectroscopy to answer the following questions: Do we observe the predicted distortions namely, flattening of spectra and the appearance of an auxiliary minimum near 230 nm at high peptide / protein concentrations? And are these changes, which point to the onset of phase separation, accompanied by a loss of helicity?

The results of our CD titrations are shown in **Fig. 4**. Data for the CC peptide show the flattening of the spectra and the appearance of a new minimum near 230 nm (**Fig. 4A**). These changes are manifest above a threshold concentration that is a precursor of phase separation. The onset of the distortion of CD spectra is also observed for the CC^-^ and CC^+^ peptides (**Fig. 4B, C**). The CD spectra for the CC^-^ peptide, which has negligible helicity, show monotonic changes as a function of peptide concentration with distorted spectra being manifest at the lowest concentrations of the three peptides studied here (**Fig. 4B**). For CC^+^, the distortion follows a loss of helicity, which is realized as the concentration goes above a threshold (**Fig. 4C**).

**Figure 4:**
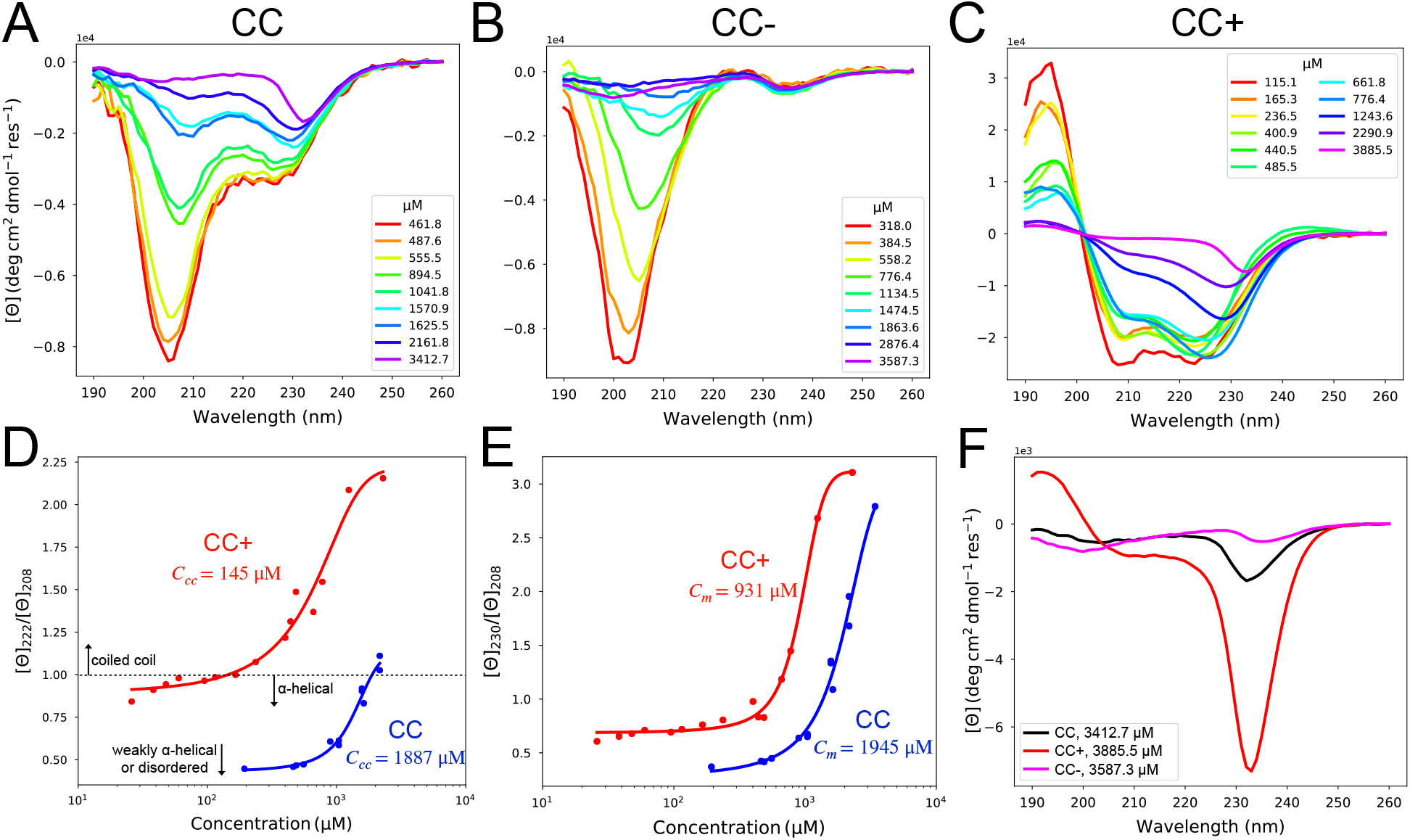
Concentration-dependent CD spectra reveal differences in conformational and assembly states across the different CC peptides. (A) CD spectra for the CC peptide. (B) CD spectra for the CC^-^ peptide. (C) CD spectra for the CC^+^ peptide. (D) The metric for coiled coil formation is shown as the ratio of the ellipticities at 222 nm and 208 nm from panels (A) and (C). The dotted line indicates when this ratio is one, above which coiled-coil formation is likely. (E) The metric for higher-order assembly is the ratio of ellipticities at 230 nm and 208 nm from panels (A) and (C). (F) CD spectra at the highest concentrations measured for each peptide.

To understand the interplay of helicity and higher-order assembly in the CC and CC^+^ peptides, we defined two metrics based on the concentration-dependent CD spectra. The ratio of ellipticities at 222 and 208 nm as a function of concentration provides a readout of coiled-coil formation and is a proxy for dimerization (64) (**Fig. 4D**). Specifically, coiled coil formation becomes more probable as the ratio of ellipticities at 222 and 208 nm exceeds one. As expected, CC^+^ forms coiled coils at much lower concentrations than CC, depicted as *c*_cc_ in **Fig. 4D**. Our second metric is defined as the ratio of the ellipticities at 230 and 208 nm as a function of concentration (**Fig. 4E**). This provides a measure of higher-order assembly, which may include bundles of helices and / or coiled coils. These data are well-fit by a 2-state model, with each peptide showing a higher *c_m_* for higher-order assembly than for coiled-coil formation. Again, CC^+^ forms higher-order assemblies at lower concentrations than CC, which is consistent with data in Fig. 3 showing enhanced dilute phase oligomerization for the corresponding full-length protein. We calculated the secondary structure propensities for the spectra from each peptide using the CDSSTR algorithm (**Fig. S4**). CC is partially helical at low to intermediate concentrations, CC^-^ demonstrates negligible helicity, while CC^+^ is strongly helical at all but the highest concentrations.

We can compare the extent to which each of the peptides drive phase separation through unfolded conformations by comparing the spectra collected at similar, high peptide concentrations (**Fig. 4F**). This comparison reveals that the extent of distortion and flattening are most pronounced for CC^-^ followed by CC, and then CC^+^. These trends are apparent despite the enhanced higher-order assembly of CC^+^, suggesting that a third level of assembly occurs in these peptides which is distinct from oligomer formation. This result is consistent with expectations that increased helicity detours peptides away from forming condensates by sequestering them into dilute phase oligomers. In contrast, the interactions among unfolded peptides appear to drive phase separation.

Finally, we asked if the observations reported for the peptides are recapitulated for the full length Whi3 protein. For this, we measured concentration dependent CD spectra for the full-length Whi3 protein (**Fig. S5**). At low concentrations, the contribution of the folded RRM is apparent in the CD spectrum. However, as concentrations increase, we observe the expected flattening and appearance of a new minimum at ~230 nm. We note that the assemblies observed in these CD experiments are precursors of condensates, since condensates themselves can become sufficiently large and fall out of solution. Our data suggest that unfolding is a pre-requisite for driving Whi3 phase separation. These data are in accord with the findings from the LASSI-based simulations, wherein folded states suppress phase separation via stronger, anisotropic interactions.

### Dense phase concentrations correlate with material states of condensates

In addition to changes in dense phase stoichiometry and dilute phase oligomer distributions, condensate morphologies in the Whi3-*CLN3* systems varied with time and across mutants, with each system showing distinct dynamical behaviors. An hour after mixing, the condensates are small and highly mobile. At intermediate stages of coarsening, droplets collide and partially coalesce (**Fig. 5A**). WT condensates have irregular shapes at the five-hour mark and later, while Whi3 (CC → CC^-^) condensates are always more spherical and round up into spheres within four hours.

**Figure 5:**
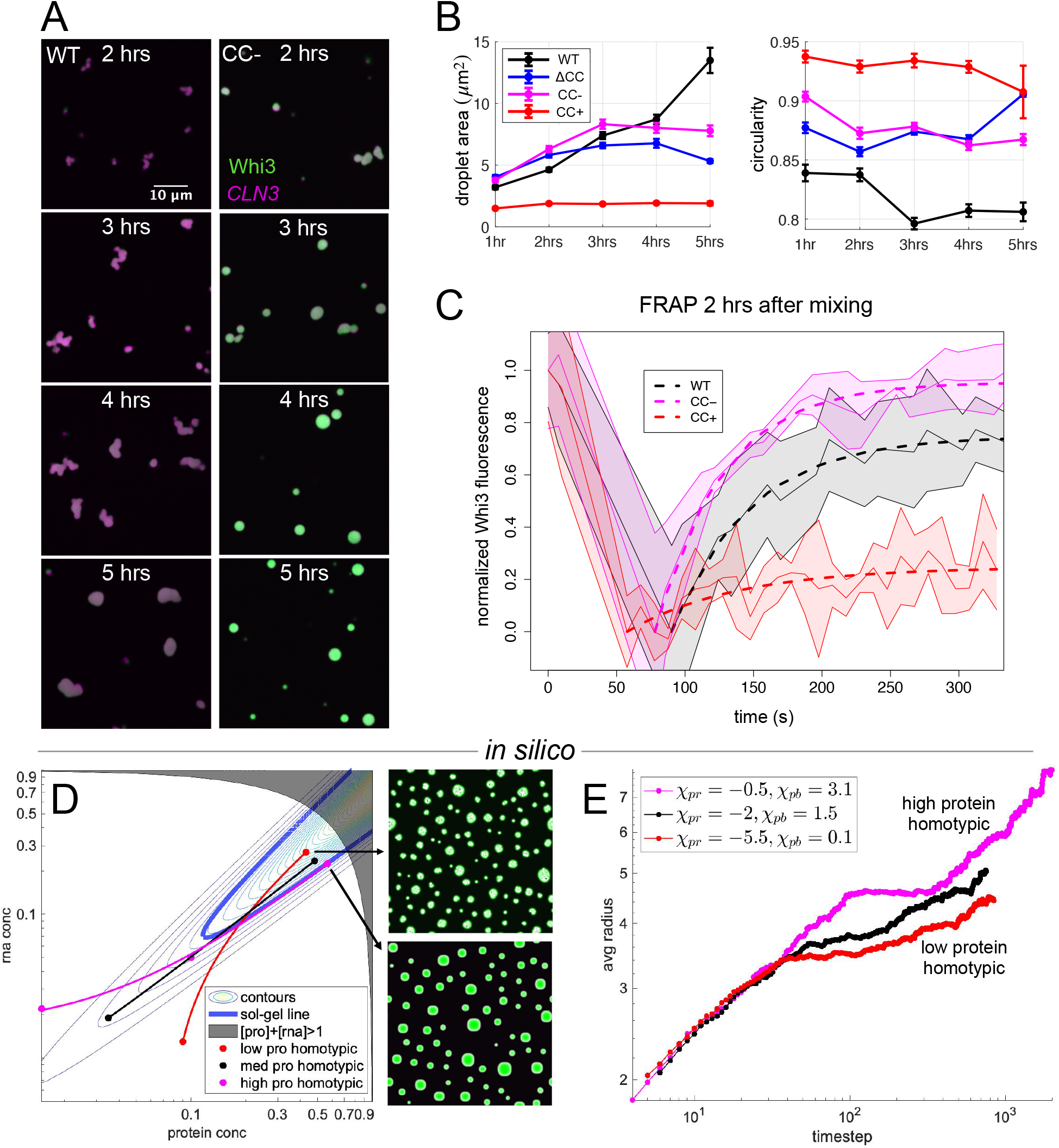
Droplet dynamics are linked to dense phase stoichiometries. (A) Representative images at each time point for WT and Whi3 (CC → CC^-^) proteins at bulk concentrations of 500 nM Whi3 with 5 nM *CLN3*. (B) Quantification of average droplet area and circularity for experiments corresponding to panel (A). Error bars are standard error of the mean. (C) FRAP curves for Whi3-*CLN3* mixtures performed 2 hours after mixing. FRAP curves for 5-8 droplets for each protein were recorded; dashed lines indicate model fits to the mean, and the shaded areas indicate +-1 standard deviation. (D) Parabolic sol-gel contours with the sol-gel line in bold blue are overlaid by dilute, bulk, and dense phase equilibrium concentrations of simulated systems with different protein homotypic interaction energies. For fixed bulk concentrations of protein and RNA, alteration of the protein homotypic interaction energies results in dense phase concentrations that lie at varying depths beyond the sol-gel line. Simulation snapshots after steady state has been reached in the low protein homotypic and high protein homotypic interaction energy systems are shown. (E) The average radii of droplets from simulations in panel (D) are shown as a function of time. The legend provides values for the interaction parameters in the Flory-Huggins free energy that determine whether the systems encode low, medium, or high protein homotypic interaction energies.

We quantified condensate shapes and sizes at different time points using a custom image analysis pipeline (see methods). The sizes of WT Whi3-*CLN3* condensates, measured in terms of their surface areas, increase as a function of time. We observe a concomitant decrease in circularity that is consistent with decreased sphericity of the condensates. Compared to WT, Whi3 (CC → CC^-^) and Whi3 (ΔCC) have larger average condensate areas at early times and always have higher circularity. This is consistent with the observation that Whi3 (CC → CC^-^) condensates coalesce more rapidly (**Fig. 5B**). The larger areas at longer time points for condensates formed by WT Whi3 and *CLN3* are partly due to the irregular shapes of these condensates, given that spheres minimize surface areas compared to other geometries.

The condensates formed by Whi3 (CC → CC^+^) with *CLN3* demonstrate distinct coarsening behavior, with very small average sizes that plateau at two hours. Interestingly, their high circularity persists throughout the time periods interrogated. (**Fig. 5B**). This is because droplets formed by Whi3 (CC → CC^+^) collide at early stages of phase separation (1-2 hrs), but instead of coalescing they either stick to one another or bounce off each other without fusing (**Fig. S6** in *SI Appendix*). Taken together, the data for different Whi3-*CLN3* condensates demonstrate a correlation between the dense phase stoichiometry on the one hand and morphologies and coarsening rates of condensates on the other. Specifically, higher Whi3-to-*CLN3* concentration ratios within condensates lead to more round droplets that coalesce more rapidly. We also observed that for a fixed concentration of 5 nM for *CLN3*, higher bulk concentrations of WT Whi3 lead to more Whi3 in dense phase droplets (**Fig. S7 A,C**). This is in line with the phase behavior that is expected for a ternary mixture of RNA, protein, and solvent (**Fig. S8A**). Condensates with more Whi3 also round up and coarsen more quickly (**Fig. S7 A,B**). These data further demonstrate a correlation between dense phase stoichiometry on the one hand and morphology and coarsening rates on the other.

We performed fluorescence recovery after photobleaching (FRAP) experiments to assess the mobilities of Whi3 and *CLN3* within condensates and to uncover how internal molecular dynamics correlate with the morphological differences noted above. Two hours after mixing 1 μM Whi3 with 5 nM *CLN3*, condensates showed differential protein mobilities that were consistent with the morphological interpretations above; WT demonstrated intermediate recovery, Whi3 (CC → CC^-^) recovered most fully and rapidly, and Whi3 (CC → CC^+^) showed minimal recovery of fluorescence after photobleaching (**Fig. 5C**). Details of experiments, data normalization, and model fits can be found in the *SI* Appendix and **Fig. S9**. In contrast to the Whi3 fluorescence, the *CLN3* fluorescence did not recover to any extent (**Fig. S10**). This is consistent with recent observations of RNA and protein molecules featuring drastically different internal mobilities in protein-RNA condensates (10, 65). We also measured FRAP on condensates formed with WT Whi3 at different bulk concentrations mixed with 5 nM *CLN3* to test whether dense phase stoichiometry alone affects internal dynamics and material properties. Indeed, as the bulk protein concentration is lowered, dense phase protein mobility decreases, although to a lesser extent than with the CC mutant systems (**Fig. S7D**). This observation indicates that there are contributions of the CC region to the interactions amongst molecules within the dense phase that further modulate protein mobility. These data confirm that *CLN3* mobility is consistently lower than Whi3 mobility and that higher Whi3-*CLN3* dense phase concentration ratios promote greater protein mobility. Differences between protein mobilities across condensates formed by different Whi3 constructs are due to differences in the driving forces for phase separation that arise from mutations to the CC region. We explore these effects by modeling the impacts of dense phase stoichiometry on the dynamics of condensate aging, as discussed next.

Our data and previous work show that Whi3-*CLN3* condensates age, resulting in slowed dynamics and fibril-like structures within droplets at long times (33). To explore the dynamics of maturation, we modeled the evolution of condensates formed by the different Whi3 variant systems using the ternary Cahn-Hilliard-Cook system of equations coupled to a gelation process (66):

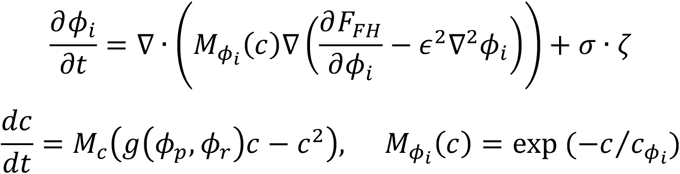

Here, *ϕ_i_* = *ϕ_p_, ϕ_r_* represent local protein and RNA volume fractions respectively, *c* represents the local gel concentration, *F_FH_* is the ternary Flory-Huggins free energy, and *g*(*ϕ_p_, ϕ_r_*) defines the parabolic sol-gel contours shown in **Fig. 5D** (derivations in *SI Appendix*). This model provides a link between dense phase stoichiometry and aging by coupling the mobility of protein and RNA, *M_ϕ_i__*. (c), to a local gel concentration whose dynamics in turn depend on the gap between the protein and RNA volume fractions and the sol-gel line. The further the system lies beyond the sol-gel line, the faster the gel concentration grows. This lowers the mobilities of protein and RNA.

At equilibrium, condensates formed by associative polymers with sticker-and-spacer architectures are proposed to form condensate spanning networks (14, 17, 18). However, the gap between the sol-gel line and the dense phase arm of the phase boundary (67) can lead to gelation without phase separation (68, 69), which corresponds to dynamically arrested phase separation. If the gap between the sol-gel line and the dense phase arm of the phase boundary is large (67), then gelation without phase separation is readily realized via dynamical arrest of phase separation (19, 67). This results in a combination of irregular morphologies and ultraslow relaxation to spherical morphologies (10, 70). Thus, the material state and resulting dynamics of condensates can be tuned by the dense phase stoichiometry and the gap between the sol-gel line and the dense phase arm (67).

We can tune dense phase stoichiometries by changing the magnitude of Whi3 homotypic interactions in the Flory-Huggins free energy (**Fig. S8B, C** in *SI Appendix*). The Flory-Huggins free energy represents averaged, non-specific interactions among components, so the Whi3 homotypic interaction in this context encompasses contributions from both the CC and QRR. The strength of the homotypic interactions that drives phase separation is implicitly linked to the degree of unfoldeness of the CC motif and the QRR in Whi3. Accordingly, in the FH model, when the Whi3 homotypic interaction strength is low, the dense phase arm is furthest beyond the sol-gel line and droplet dynamics are readily arrested, as in the Whi3 (CC → CC^+^) system (**Fig. 5 D, E**). Alternatively, when the Whi3 homotypic interactions strength is high, the gap between the dense phase concentration and sol-gel line is narrowed, and the dynamics are minimally affected (**Fig. 5 D, E**). We observed similar effects when altering only Whi3 bulk concentration in the model and fixing interaction energies. Here, high Whi3 bulk concentrations lead to high dense phase Whi3 concentrations and coarsening over longer times (**Fig. S7 E, F**, *SI Appendix*). These results are a natural consequence of a ternary phase boundary intersecting the sol-gel line, and they represent the influence of stoichiometry on gelation.

Since the model does not include Brownian motion of droplets, coalescence does not contribute to coarsening. Instead, coarsening is driven by Ostwald ripening (71), which is hindered as gelation arrests phase separation. Thus, droplet circularity is not affected by gelation in this model, but droplet sizes are affected, whereby smaller sizes indicate increased gelation that arrests phase separation. We confirmed this by performing simulations at different homotypic interaction strengths and bulk concentrations without the gelation term, an approximation that corresponds to the scenario of phase separation without gelation. In this scenario, we do not find significant differences in average droplet sizes (**Fig. S11**, *SI Appendix*). Overall, the results from the modeling are consistent with the experimental observations that a higher dense phase Whi3 to *CLN3* concentration ratio correlates with increased mobility of proteins within condensates. The modeling also highlights the importance of dense phase stoichiometries in impacting the sequencespecific gaps between sol-gel lines and dense phase arms of phase boundaries (67).

## Discussion

We have identified a 21-residue motif, designated as CC, that lies within the Whi3 QRR. In WT Whi3, the CC motif is likely defined by an equilibrium between helical and non-helical conformations. Mutations that alter the helicity of this motif can be used to tilt the equilibrium toward conformations that are more helical or less helical. Altering the helical propensity of CC through mutations results in trade-offs of order and disorder at the molecular level. Interestingly, these differences have profound effects on the phase behavior of the Whi3-*CLN3* system. Interactions among Whi3 molecules with stable helical CC motifs lead to a sequestration of proteins into dilute phase oligomers. This leads to a lowering of protein concentrations in the dense phases of RNP condensates. It also contributes to a faster aging of RNP condensates because the gap between the sol-gel line and dense phase arm of the phase boundary is widened. In contrast, increased disorder within the CC motif and disorder within the remainder of the QRR weakens dilute phase oligomerization and drives Whi3 molecules into RNP condensates. These condensates experience slower aging due to a minimized gap between the sol-gel line and the dense phase arm of the phase boundary.

Interestingly, we find that the Whi3 (ΔCC) construct behaved most like the Whi3 (CC → CC^-^) construct. Specifically, we observe higher protein concentrations in the dense phase for Whi3 (ΔCC) when compared to WT at similar bulk concentrations of each protein and hence lower bulk volume fractions of Whi3 (ΔCC) compared to WT. Clearly, the effect of removing the CC is strong enough to compensate for a small length difference that should weaken phase separation of Whi3 (ΔCC). Deletion of the CC motif would imply that the QRR seldom samples helical conformations, and phase separation is driven via interactions among disordered regions. For WT, we hypothesize that helicity from the CC is sufficient to tip the balance toward interactions among helical domains that sequester Whi3 proteins into dilute phase oligomers.

Interactions among Whi3 and *CLN3* drive phase separation, primarily through heterotypic interactions that involve a disordered QRR and a folded RRM. Homotypic interactions mediated by oligomerization via short helical domains detour the molecules away from co-phase separation with RNA. The LASSI simulations suggest that the valence of stickers within the QRR contribute to homotypic interactions, which are potentially weakened by competing heterotypic interactions with *CLN3* in the ternary system as indicated by lower dense phase protein concentrations in the ternary system (**Fig. 2B**) relative to those in the binary system (**Fig. 2D**). The question is why do we observe clusters that span a range of sizes in the dilute phase (**Fig. 3C,D**)? The clusters are likely to involve a combination of stoichiometric assemblies mediated mainly by interactions among helical motifs, and heterogeneous distributions of clusters that are mediated by the CC motif and the QRR. These heterogeneous distributions are manifestations of PSAP, whereby pre-percolation clusters of different sizes are expected to be present in the sol that coexists with the dense phase (16, 19, 72).

The sequence motifs we have identified within the Whi3 QRR are common features of proteins with QRRs. Indeed, other groups have shown that Leu, Pro, and His are especially enriched within and around QRRs, with Leu being typically clustered at N-terminal regions and Pro being prevalent at C-terminal regions (73, 74). Here, we observe similar positioning of these residues within the Whi3 QRR (**Fig. 1A**). In accord with our findings, Kokona and Fairman have shown that formation of stable coiled coils suppresses polyQ driven insolubility via aggregation and the formation of solid-like inclusions by peptides that mimic the exon 1 encoded region of huntingtin (75). Interestingly, Ford and Fioriti have demonstrated that QRRs, coiled-coils, and RNA-binding domains commonly co-occur, especially in neuronal proteins that are implicated in both phase separation and neurogenerative diseases caused by toxic, amyloid assemblies (76).

### Relation to other systems with oligomerization domains and IDRs

Many proteins that have been identified as drivers of phase separation contain both structured oligomerization domains and IDRs. Reports to date have shown that oligomerization promotes phase separation (2–4, 12, 37–39, 49, 77). A notable exception to these findings is the prediction of a “magic-number effect” that occurs for rigid domains with sufficiently strong, specific interactions, whereby molecules become trapped in stoichiometric assemblies (78). Similar effects are realizable, irrespective of the strength of inter-domain interactions, if the linkers between the interacting domains are too short (17). In the model that shows the magic number effect, binding partners with an integer multiple of one-to-one binding sites can saturate interactions with each other, favoring the formation of network terminating oligomers in the dilute phase. The experiments we summarize here show the phenomenon of dilute phase oligomers sequestering the Whi3 protein out of the dense phase, although this behavior does not appear to be a magic-number effect. It is possible that the CC motif may prefer a specific oligomerization state, which on its own could result in stable oligomers with saturated binding sites. However, Whi3 also mediates homotypic interactions through the rest of its QRR that presumably has no fixed valence. It also participates in heterotypic interactions with RNA via its RRM. These interactions contribute complexity to the system that makes it difficult to interpret in the context of a magic number effect, although the phenomenological similarity between the theory of Xu et al. (78) and the effects of increasing the CC stability (rigidity) and interaction energy are intriguing.

The presence of structural motifs embedded within IDRs may provide an explanation for the effects of disease-related point mutations, as has been shown for TDP-43 (49). If such mutations perturb structural motifs and their associations, then drastic shifts in phase behavior and material states are possible even with point mutations (79). Importantly, the effects of the CC motif that we have uncovered in this study are distinct from those in TDP-43, highlighting the need for context-dependent studies of embedded structured domains. Identification of such motifs within phase separating systems will be an essential task for improved understanding of the driving forces that contribute to the formation of biomolecular condensates.

## Materials and Methods

### UV-Circular Dichroism

Peptides were purchased in pure form from GenScript with N-terminal acetylation and C-terminal amidation. All peptide sequences included an N-terminal tryptophan to enable concentration measurements, followed by a glycine, and then the sequence of interest. CD measurements were carried out using a Jasco 810 spectropolarimeter scanning from 260 nm to 190 nm, with a data pitch of 1 nm and a bandwidth of 1 nm. Four to six accumulations were averaged for each spectrum with a scanning speed of 50 nm/min and a two second response time. Additional details regarding the CD measurements are described in the *SI Appendix*.

### Assembly and imaging of *Whi3-CLN3* condensates *in vitro*

Labeled and unlabeled stocks of Whi3 protein (WT, CC+, CC-, ΔCC) and labeled *CLN3* mRNA were stored in 200 μL, 33 μL, and 10 μL aliquots respectively at −80°C until just before experiments. Aliquots were thawed on ice and protein samples were spun down at 13,200 rpm for 5 minutes to remove any aggregates. The concentration of labeled and unlabeled protein was then determined by measuring absorbance at 280 nm. The labeled protein concentration was corrected using a factor of 0.09 to account for the Atto 488 absorbance, and labeling efficiency was determined as the ratio of the dye concentration to the corrected protein concentration. Labeled and unlabeled Whi3 proteins and *CLN3* mRNA were diluted into buffer containing 50 mM HEPES at pH 7.4 and 150 mM KCl to desired concentrations and labeled fractions of protein and mRNA. The volumes of labeled and unlabeled protein stock needed to simultaneously obtain a desired total protein concentration and labeled fraction were determined using the following formulas: 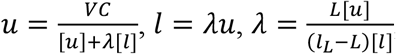, where *u* is the desired volume of unlabeled protein (μL), *l* is the desired volume of labeled protein (μL), *V* is the final reaction volume (200 μL for phase separation assays, 50 μL for FRAP assays), *C* is the desired final protein concentration (μM), [*u*] is the measured concentration of the unlabeled protein stock (μM), [*l*] is the measured concentration of the labeled protein stock (μM), *l_L_* is the measured labeled fraction of the labeled protein stock, and *L* is the desired protein labeled fraction (5% for phase separation assays, 10% for WT and CC^-^ and 20% for CC^+^ in FRAP assays). Whi3 proteins were prepared at three concentrations, 250 nM, 500 nM, and 1 μM. *CLN3* mRNA was prepared at a concentration of 5nM for every condition. For phase separation assays, 200 μL of protein and mRNA solutions were loaded into glass-bottom imaging chambers (Grace Bio-Labs) that were pre-coated with 30 mg/mL BSA (Sigma) for 30 mins to prevent protein adsorption to the surfaces of the well. The surfaces were washed thoroughly with buffer prior to addition of protein and mRNA solutions. Labeled and unlabeled Whi3 stocks were added first to imaging wells, followed by *CLN3*, and the final solution was mixed by pipetting without introducing bubbles. Imaging chambers were either placed in a 25C incubator until imaging or immediately transitioned to a temperature-controlled microscope stage (Tokai Hit from Incubation System for Microscopes) set to 25C. For each desired timepoint, confocal z-stacks of imaging wells were acquired. Three separate z-stacks within each well were taken at each time point at random locations. Fluorescence images in two channels were acquired using 488 nm and 561 nm lasers to visualize Atto488-labeled Whi3 and Cy3-labeled *CLN3* respectively. An Atto 488 dye calibration curve was created by imaging 200 μL wells with varying concentrations of dye. Experiments were replicated three times.

### LASSI simulations

Simulations were performed using LASSI and run on the Longleaf computing cluster at UNC-Chapel Hill. Each simulation was run independently on a single compute node with 4GB RAM. Interaction energies that were used for each of the three systems studied (WT, CC+, CC-), are shown in **Table S2** *SI Appendix*.

## ACKNOWLEDGMENTS

This work was supported by grants from the US National Institutes of Health (5R01NS056114 and R01NS089932 to RVP, R01BM081506 to ASG, F32-GM133123-01A1 to BMS, F32-GM136055 to WTS, and T32-GM8570-25 to IS), the Air Force Office of Scientific Research (FA9550-20-1-0241 to ASG and RVP), the HHMI faculty scholars (to ASG). We are grateful to Furqan Dar and Kiersten Ruff for helpful discussions. Furqan Dar provided guidance and assistance with LASSI simulations.

