## Supplementary material for "Dilute phase oligomerization can oppose phase separation and modulate material properties of a ribonucleoprotein condensate": SI Appendix

#### This PDF file includes:

Supplementary text
Figs. S1 to S11 (not allowed for Brief Reports)
Tables S1 to S2 (not allowed for Brief Reports)
SI References

### Supporting Information Text

**UV-CD measurements.** A demountable quartz cuvette with a path length of 0.1 mm was used for all measurements. The buffer used in all measurements was 20 mM HEPES, pH 7.4 with 150 mM NaF, filtered with a 0.2 $\mu$m PES filter. Fluoride was used in place of chloride as the anion to avoid the signal interference caused by chloride. For concentration-dependent CD measurements, a high concentration stock (3-4 mM peptide) was prepared by dissolving lyophilized peptide in a small volume of buffer. Subsequent samples were obtained by dilution of the stock solution using the same buffer. For CD measurements acquired as a function of temperature, the temperature was varied from 5°C to 95°C or from 95°C to 5°C at a rate of 5°C per minute, with a one minute pause before each full spectrum measurement made at 10°C intervals. Single wavelength (either 208 or 222 nm) measurements were made at 5°C intervals. Peptide concentrations were prepared in the 600-900 $\mu$M range. CD spectra were converted from machine units (mdeg) to mean residue ellipticity ($[\theta]$, deg cm$^2$ dmol$^{-1}$ residue$^{-1}$) using the equation $[\theta] = \frac{\theta}{(N-1)LM}$, where $\theta$ is machine units, $N$ is the number of amino acids in the peptide, $L$ is path length and $M$ is molar concentration. Peptide concentrations were determined by measuring absorbance spectra using an Implen NP80 nanophotometer. Molar concentration values were calculated using an extinction coefficient of 5500 M$^{-1}$ cm$^{-1}$ at 280 nm.

In **Fig S1**, mean residue ellipticity at 222nm was fit for each peptide with the following 2-state model:

$$[\theta]_{222nm} = \frac{A_1 - A_2}{1 + \exp\left(m(T_m - T)\right)} + A_2 \quad [1]$$

**Deconvolution of CD data.** CD Data sets were converted to units of mean residue ellipticity as described above and submitted to the Dichroweb server (1) (http://dichroweb.cryst.bbk.ac.uk/html/home.shtml) for deconvolution using the CDSSTR algorithm (2) and basis sets 4, 7 (3), and SP175 (4). These are the basis sets that have been optimized for the wavelength rage of 190-240 nm. We explored algorithms other than CDSSTR, including CONTIN and SELCON3 (5), but CDSSTR consistently provided the best fits for our data, so the other algorithms were not used further. We note that the output of the analysis is broken down into two helical categories and two beta strand categories (6), but for the sake of simplicity we present the results in terms of a single helical estimate and a single beta strand estimate, each of which is the sum of the respective sub-categories. Reported secondary structure content estimates represent the mean and standard deviation of the values resulting from analysis using each of the three basis sets (4, 7, SP175).

**Bacterial expression and purification.** Whi3 constructs were transformed into BL21 competent cells (New England Biolab C2527I). Cells were grown at 37C with shaking and appropriate selection until an OD600 of 0.6 was reached. Then cells were shifted to 18C and induced with 1mM IPTG (Fisher I2481C25). Cells were grown for 18hr and then pelleted by centrifugation at 13000rcf. The Pellet was resuspended in 25ml Lysis buffer (1.5M KCl, 20mM Imidazole, 5mM beta-mercaptoethanol, 50mM Hepes pH7.4) along with 1 tablet Pierce Protease inhibitor (Thermo-FIsher Scientific A32965) and 15mg Lysozyme (Fisher BP535-10). Cells were lysed at 4C with gentle rocking for 1hr and then briefly sonicated on ice. Lysate was clarified by centrifugation at 50000rcf for 30min. Clarified lysate was mixed with 0.8mL of Cobalt resin (Thermo Fisher Scientific PI89965), column was washed with 5 column volumes of lysis buffer and then eluted with 3ml of elution buffer (150mM KCl, 200mM Imidazole, 5mM beta-mercaptoethanol, 50mM Hepes pH7.4). Protein was dialyzed into storage buffer (150mM KCl, 50mM Hepes pH 7.4) using a slide-a-lyzer cassete and a 20,000MW cutoff (Thermo-Fisher Scientific 87735). Protein was labelled using an atto-488-NHS ester (Sigma Aldrich 41698) and then dialyzed again to remove uncongugated dye.

**RNA *in vitro* transcription.** Plasmid was linearized with XbaI and NotI restriction enzyme (New England Biolabs R0145S, R0189S) and gel purified (QIAGEN 28706). 100ng of gel purified DNA was used as a template for in vitro transcription (NEB E2040S) carried out according to the manufacturer's instructions with the addition of Cy3 (Sigma PA53026) labeled UTP to each reaction. Following incubation at 37C for 18 hours, in vitro transcription reactions were treated with DNAseI (NEB M0303L) according to the manufacturer's instructions. Following DNAse treatment, reactions were purified with 2.5M LiCL precipitation. Purified RNA concentrations were quantified using nanodrop and verified for purity and size using a denaturing agarose gel and Millenium RNA ladder (ThermoFisher Scientific AM7151).

**Assembly and imaging of Whi3-only condensates *in vitro*.** Whi3 proteins were concentrated using 30 kDa molecular weight cutoff centrifugal filters (Amicon) and diluted into 50 mM HEPES pH 7.4 and 2 mM TCEP buffer at the specified protein concentrations such that the final KCl concentration was 75 mM. Atto488-labeled protein was included such that approximately 4-6% of protein was dye-labeled. Protein solutions were loaded into glass-bottom imaging chambers (Grace Bio-Labs). Wells were pre-coated with 30 mg/mL BSA (Sigma) to prevent protein adsorption to the well surfaces, and washed thoroughly with experiment buffer prior to addition of protein solutions. Wells were incubated for 2.5 h at room temperature to allow droplets to form and sediment onto the surface of the imaging chambers. Confocal z-stacks of droplets were acquired, starting below the coverslip surface, with a z-spacing of 0.2 $\mu$m. Imaging was repeated 4 h post-mixing. At least five z-stacks of each condition were acquired, and all experiments were replicated three times per condition.

**Fluorescence recovery after photobleaching (FRAP).** Silicone gaskets were laid onto plasma-cleaned coverglass and served as the reaction compartments for imaging of Whi3-*CLN3* droplets. Reaction wells were passivated with BSA for 15 mins and then washed thoroughly with buffer (150 mM KCl, 50 mM HEPES pH 7.4, 5 mM BME) before adding buffer, Whi3, and *CLN3*. 50 $\mu$L volumes were used in each well, and components were assembled as described in the section: Assembly and imaging of Whi3-*CLN3* droplets *in vitro* in the Methods section of the main text. 1 $\mu$M Whi3 and 5 nM *CLN3* were used for all experiments. The coverglass was placed in a petri dish to prevent evaporation and incubated at 25C for 2 hours before imaging. During imaging, 4-8 droplets were photobleached for each experiment, and a similar number of similarly-sized droplets in each field of view were left unbleached for use in later correction of bleaching due to imaging. The data were processed in FIJI using the Time Series Analyzer which allows for selection of ROIs corresponding to both bleached and unbleached droplets. The following analysis was carried out for each of the Whi3 and *CLN3* channels. The fluorescence signals for bleached and unbleached droplets were averaged separately to obtain the following average fluorescence intensities as a function of time: $\hat{F}(t)$, the averaged bleached droplet fluorescence, and $\hat{B}(t)$, the average unbleached droplet fluorescence. A corrected signal was calculated by dividing the bleached droplet signal at each timepoint by the unbleached droplet signal: $\hat{F}_{corrected}(t) = \frac{\hat{F}(t)}{\hat{B}(t)}$. The fluorescence before bleaching is defined as $F(0) = \hat{F}_{corrected}(t = 0)$, and the fluorescence just after bleaching is defined as $F_0 = \hat{F}_{corrected}(t = t_{bleach})$. We normalized $\hat{F}_{corrected}(t)$ by $F(0)$ and $F_0$ using the following formula: $N(t) = \frac{\hat{F}_{corrected}(t) - F_0}{F(0) - F_0}$. We fit $N(t)$ to the following formula: $N(t) = A(1 - \exp(\frac{-t}{\tau}))$. We calculated the standard deviation of the original fluorescence curves, sd($F(t)$), and propagated the error through the normalization process. This amounts to dividing the standard deviation at each time by the factor $\hat{B}(t)(F(0) - F_0)$. $N(t)$ plus and minus the propagated standard deviation and with the model fit overlaid is shown in the main text in **Fig 5C**, and the raw data with normalizations and model fit parameters are shown in **Fig S9**.

**Calculating protein concentration from fluorescence calibration curves.** We used dye solution standards to determine protein concentrations. Briefly, dye solution standards were created by diluting Atto488 dye to concentrations ranging from 0-10 or 0-20 $\mu$M in the same buffer used for droplet assembly. Solutions were loaded into imaging wells and at least three confocal z-stacks of dye standards, starting below the coverslip surface, were acquired using the same laser power and exposure settings used for droplet imaging. For Whi3-only phase separation assays (**Fig 2C,D**, two different laser settings were used for WT and CC mutant droplets, hence the two plots in **Fig S2B**. Dye intensity was determined by computing the mean intensity of the dye image frame with the highest intensity value. For experiments including both Whi3 and *CLN3*, all proteins were labeled at 5%. For Whi3-only phase separation assays, estimated labeled populations of the constructs were WT: 5.95%, CC-: 4.15%, and CC+: 4.28%. Fluorescence values in condensates are converted to dye concentrations using the fits in **Fig S2** and divided by labeled population for each construct to estimate protein concentrations.

**Analysis of *in vitro* condensates.** Images were analyzed using a custom FIJI macro and the output data was further analyzed in Matlab to perform statistics and plotting. For each image, the FIJI macro loops through each z-slice in the mRNA channel for Whi3-*CLN3* experiments, and the single Whi3 channel for Whi3-only experiments. The mRNA channel is used for Whi3-*CLN3* experiments because the mRNA intensity is constant across experimental conditions, while the protein fluorescence intensity varies widely. For each slice, the image is duplicated, and on the duplicate a Yen threshold (7) on the fluorescence intensity is performed, and a watershed operation (8) is run on the resulting binary image. Using masks from the binary image, the "Analyze Particles" operation within FIJI is run on the original image and records the area, mean fluorescence intensity, and circularity for objects larger than 50 pixels in area. The area of all detected objects is summed and recorded. The slice with the largest total area for each image is chosen as the optimal slice, and statistics in both *CLN3* and Whi3 channels are recorded and used for further analysis. For Whi3-only experiments, only the single channel is used.

In Matlab, Whi3 fluorescence intensities are converted to concentrations using the calibration curves. Data from three imaging frames per replicate per experimental condition at each time point were aggregated across three replicates. Data are plotted either as boxplots at single time points, or as line plots with means plus and minus the standard error of the mean over time.

**Determination of proteins per puncta on passivated surfaces.** Glass coverslips were passivated with polyethylene glycol (PEG) by coating the glass with a layer of poly-L-lysine (PLL) conjugated to PEG. PLL-PEG was made according to a previous protocol (9). Briefly, amine-reactive PEG-SVA (succinimidyl valerate) was added to a 40 mg/mL mixture of PLL in 50 mM sodium tetraborate pH 8.5 at a molar ratio of one PEG per five lysine subunits. The mixture was stirred continuously for 6 h at room temperature and buffer exchanged into 50 mM HEPES pH 7.4 and 150 mM KCl using Centri-Spin size exclusion columns (Princeton Separations). Imaging wells were made by placing silicone gaskets onto oxygen plasma-treated coverslips. Wells were coated for 20-30 min with PLL-PEG diluted tenfold in 50 mM HEPES pH 7.4 and 75 mM KCl buffer. After coating, the well was washed repeatedly with HEPES buffer to remove excess PLL-PEG. Whi3 proteins were then diluted in 50 mM HEPES pH 7.4 and 75 mM KCl buffer and added to PEG-coated coverslips at concentrations of 100 or 150 nM. Proteins were allowed to settle on the surface for approximately 10-20 min, and images of puncta were acquired using TIRF microscopy.

Images of puncta were cropped to the center 400x400 pixels (center 1/9) of the original images to ensure even illumination across the field of view. Diffraction-limited protein puncta were detected using cmeAnalysis particle detection software (10), which fit puncta to a two-dimensional Gaussian function with standard deviation, $\sigma$, determined from the TIRF microscope point-spread function (PSF).

Briefly, the PSF $\sigma$ was determined by acquiring images of 100 nm TetraSpeck fluorescent microspheres (Invitrogen) on a coverslip surface using the same TIRF settings used in experiments. The fluorescence intensity profiles of diffraction-limited microspheres were fitted to the two-dimensional Gaussian function using Wolfram Mathematica 12 software. The PSF $\sigma$ was taken as the average from seven fits.

The cmeAnalysis software reports the amplitude, "A," of the Gaussian fit over the local background intensity, "c." The A values were accepted as valid if they came from diffraction-limited puncta and were significantly above the local c values. The number of proteins per puncta was computed from A values by comparing to the intensity of single protein molecules, described in the next section.

**Determining the intensity of single protein molecules.** Silicone gaskets were placed on oxygen plasma-treated coverslips and Atto488-labeled Whi3 was added at a concentration of 50-100 pM. Images of single protein molecules adhered to the coverslip surface were acquired with TIRF microscopy using the same TIRF and laser power settings used in experiments. Images were cropped to the center 400x400 pixels of the original images and the diffraction-limited puncta of single proteins were detected using cmeAnalysis software, following a similar procedure described in the previous section. "A" values were pooled from 5-10 image frames and binned into a histogram, yielding a distribution with one clear peak corresponding to the average intensity of a single protein molecule. This value was then used to calculate the number of dye-labeled proteins per puncta. Finally, the total number of proteins per puncta was determined by adjusting for the known fraction of dye-labeled to unlabeled protein in the experiment.

Single-step photobleaching experiments were performed to validate our approach for determining single protein molecule intensity. Specifically, we prepared samples of dilute, glass surface-adhered proteins as described above and acquired time-lapse movies of protein bleaching using TIRF microscopy. The intensities of puncta over time, after background subtraction, were analyzed using ImageJ software. This approach revealed that proteins frequently bleached to the level of the local background in a single, clear step. We found that the initial, average intensities of such puncta were in agreement with particle detection-based estimates of single molecule intensity.

**Additional details regarding the LASSI simulations.** We used the open-source version that we have made available on GitHub (https://github.com/Pappulab/LASSI). We added analysis routines to enable the calculation of specific quantities that are unique to the current work.

Each chain consists of a single CC bead connected to four QRR beads, with a correspondence of approximately 20 amino acids per bead. 2000 chains were used in each simulation. For each system, 35 volume fractions were run, logarithmically spaced between volume fractions of 0.0001 to 0.79. For each volume fraction, 35 temperatures were run, linearly spaced between 1 and 2 (arbitrary units). Temperature scales interaction energies as $\frac{\epsilon}{T}$. For each temperature and volume fraction, each simulation ran for 1e9 steps, after 5e6 steps of thermalization during which the temperature is set to 1000 and polymers are forced to the center of the simulation box. Two replicates are run for each simulation. As described in (11), the last half of trajectories for each simulation across the two runs are analyzed and averaged to produce a global density inhomogeneity metric, $\bar{\rho}$. Above $\bar{\rho} = 0.025$, the system is phase separated, as detailed in (11). Binodals in this work are drawn along contours corresponding to $\bar{\rho} = 0.025$.

Dilute phase oligomer distributions were calculated with custom scripts that used the ovito python module (12). The final frame of each trajectory corresponding to a single run for each simulation condition was used for analysis. A cluster analysis was performed on the particle positions, where clusters within 2 units of each other are considered to be in the same cluster (this allows for all possible adjacent lattice positions with a 3D lattice). If the system was phase separated ($\bar{\rho} > 0.025$), the largest cluster, which corresponds to the dense phase, is removed from further analysis. The proportion of chains that are monomeric in the resulting dilute phase is calculated as the total number of chains not in a cluster divided by the total number chains in the dilute phase. Plots were created using matplotlib in python.

**Computing phase diagrams from the Flory-Huggins free energy.** To begin to understand the effects of homotypic interactions on phase behavior, we studied the Flory-Huggins free energy, a commonly used mean-field model of phase separation in polymer solutions (13). Our system consists of two polymers, protein and RNA, and the buffer, so it is characterized by the ternary form of the free energy:

$$F_{\mathrm{FH}}(\phi_p, \phi_r) = \frac{\phi_p}{N_p}\ln\phi_p + \frac{\phi_r}{N_r}\ln\phi_r + (1-\phi_p-\phi_r)\ln(1-\phi_p-\phi_r) + \chi_{pr}\phi_p\phi_r + \chi_{pb}\phi_p(1-\phi_p-\phi_r) + \chi_{rb}\phi_r(1-\phi_p-\phi_r) \qquad [2]$$

where $\phi_p, \phi_r$ are the concentrations of protein and RNA, respectively, and the third component, buffer, is expressed as $\phi_b = 1 - \phi_p - \phi_r$ due to the conservation of mass. Each interaction term is defined as: $\chi_{ij} \sim \epsilon_{ij} - \dfrac{\epsilon_{ii} + \epsilon_{jj}}{2}$, where $\epsilon_{ij}$ denotes the interaction energy between species $i$ and $j$, and is negative for attractive forces by convention. We compute phase diagrams for the system by finding dense and dilute phase concentrations that result in equivalent chemical potentials. For a given set of parameters, the system will phase separate if the chemical potentials of the dense and dilute phases are equal for each component, i.e.:

$$\left.\frac{\partial F_{\mathrm{FH}}}{\partial \phi_p}\right|_{\phi_p=\phi_p^L} = \left.\frac{\partial F_{\mathrm{FH}}}{\partial \phi_p}\right|_{\phi_p=\phi_p^H}$$
$$\left.\frac{\partial F_{\mathrm{FH}}}{\partial \phi_r}\right|_{\phi_r=\phi_r^L} = \left.\frac{\partial F_{\mathrm{FH}}}{\partial \phi_r}\right|_{\phi_r=\phi_r^H} \quad [3]$$
$$\left.\frac{\partial F_{\mathrm{FH}}}{\partial \phi_b}\right|_{\phi_b=\phi_b^L} = \left.\frac{\partial F_{\mathrm{FH}}}{\partial \phi_b}\right|_{\phi_b=\phi_b^H}$$

Using the conservation of mass relation from above, this system simplifies to a system of three equations with the unknowns $\phi_p^L, \phi_p^H, \phi_r^L, \phi_r^H$. For a fixed value of $\phi_p^L$, for example, we can solve the system by minimizing the following residual function (14):

$$R = \sum_i \left( \left.\frac{\partial F_{\mathrm{FH}}}{\partial \phi_i}\right|_{\phi_i=\phi_i^L} - \left.\frac{\partial F_{\mathrm{FH}}}{\partial \phi_i}\right|_{\phi_i=\phi_i^H} \right)^2 \quad [4]$$

For a given set of parameters, we solve for phase diagrams using a custom global minimization routine on the residual function defined above in Matlab.

In ternary systems, two species will phase separate together out of the third if their heterotypic interactions dominate over each of their homotypic interactions. We are interested in protein and rna condensing together, so the above requirement is on their interaction term, $\chi_{pr}$. The constraint that heterotypic interactions dominate requires that $|\epsilon_{pr}| > \left|\frac{\epsilon_{pp}+\epsilon_{rr}}{2}\right|$ which implies that $\chi_{pr} < 0$. We indeed find that this is a necessary condition to produce closed loop phase diagrams. We investigated the effects of altering protein and rna homotypic interactions by fixing all other parameters and setting $\chi_{pb}$ and $\chi_{rb}$ accordingly (**Fig S5**). When homotypic interactions are equal, the phase diagram is symmetric about the line at which protein and RNA concentrations are equal (**Fig S5A**). As either protein or rna homotypic interactions are increased, the phase diagram lies more closely along that axis and its tie lines become more aligned with that axis (**Fig S5B, C**). It is evident that one can increase the dense phase concentration of a component in a ternary system by either increasing the bulk concentration of that component, or by increasing the homotypic interaction energy of that component.

**Deriving the ternary Cahn-Hilliard-Cook equation coupled to gelation.** Since our system comprises protein, RNA, and buffer, a model well-suited to study the dynamics of phase separation is the Cahn-Hilliard-Cook system of equations:

$$\frac{\partial \phi_i}{\partial t} = \nabla \cdot \left( M(\boldsymbol{\phi}) \nabla \frac{\partial F}{\partial \phi_i} \right) + \sigma \cdot \zeta \quad [5]$$

where $\phi_i = \phi_p, \phi_r$, $M(\boldsymbol{\phi})$ is a concentration dependent mobility, $\zeta$ is a mean-zero Gaussian noise with variance $\sigma$, and $\frac{\partial F}{\partial \phi_i}$ is the functional derivative of the following free energy functional:

$$F(\boldsymbol{\phi}, \nabla\boldsymbol{\phi}) = \int_\Omega \big( F_{\mathrm{FH}}(\boldsymbol{\phi}) + \nabla\boldsymbol{\phi}^T \Gamma \nabla\boldsymbol{\phi} \big) d\Omega \quad [6]$$

where $F_{FH}$ is the ternary Flory-Huggins free energy. We assume that $\Gamma$ is a diagonal matrix of constants, $\epsilon^2$, for simplicity. Since we are only interested in 2-phase behavior in which protein and RNA condense together, their cross gradient terms should have either neutral or negative energies, i.e. there should be no interfaces formed between RNA and protein. To simplify the model, we assume that this energy is 0 such that all interactions are encoded in the $\chi_{ij}$. Taking the functional derivative of $F$, we arrive at the following form:

$$\frac{\partial \phi_i}{\partial t} = \nabla \cdot \left( M(\phi_i) \nabla \left( \frac{\partial F_{\mathrm{FH}}}{\partial \phi_i} - \epsilon^2 \nabla^2 \phi_i \right) \right) + \sigma \cdot \zeta \quad [7]$$

We introduce a gelation process to the above system of equations as described in the text. To define the log-parabolic sol-gel contours, $g(\phi_p, \phi_r)$, we have the following:

$$g(\phi_p, \phi_r) = \frac{r' \left(\frac{p'}{p_0}\right)^{-w \log(p'/p_0)} - r_0}{\max\limits_{p+r=1} g}$$
$$p' = p_0 \left(\frac{p}{p_0}\right)^{\cos(\theta)} \Big/ \left(\frac{r}{r_0}\right)^{-\sin(\theta)} \quad [8]$$
$$r' = r_0 \left(\frac{p'}{p_0}\right)^{\sin(\theta)} \Big/ \left(\frac{r}{r_0}\right)^{\cos(\theta)}$$

where $\theta$ defines the rotation angle in log-log space of the function.

**Numerical solution of the Cahn-Hilliard-Cook equation coupled to gelation.** The introduction of the gelation coupling to the Cahn-Hilliard equation introduces a mobility term which varies in time and space, thus making the entire equation non-linear. For this reason, we use the scheme first developed in (15). In the Fourier domain, our system of equations becomes:

$$\frac{\partial \tilde{\phi}_i}{\partial t} = ik \cdot \Big( M(c) \Big[ ik' \Big( \frac{\partial \tilde{F}_{FH}}{\partial \phi_i} + \epsilon^2 k^2 \tilde{\phi}_i \Big) \Big]_r \Big)_k \qquad [9]$$

where the subscript $i = p, r$ denotes the protein and rna concentrations, respectively, $F_{FH}$ is the ternary Flory-Huggins free energy, the tilde and the $k$ subscript denote the Fourier transform, and the $r$ subscript denotes the inverse Fourier transform. We stabilize the equation by adding the following linear fourth-order term to both sides: $A\epsilon^2 k^4 \tilde{\phi}_i$, where $A = \frac{1}{2}\max(\chi_{ij})$. Discretizing in time, we now have:

$$\frac{\tilde{\phi}_i^{\,t+1} - \tilde{\phi}_i^{\,t}}{\Delta t} + A\epsilon^2 k^4 \tilde{\phi}_i^{\,t+1} = A\epsilon^2 k^4 \tilde{\phi}_i^{\,t} + ik \cdot \Big( M(c^t) \Big[ ik' \Big( \frac{\partial \tilde{F}^t_{FH}}{\partial \phi_i} + \epsilon^2 k^2 \tilde{\phi}_i^{\,t} \Big) \Big]_r \Big)_k \qquad [10]$$

Solving for $\tilde{\phi}_i^{\,t+1}$,

$$\tilde{\phi}_i^{\,t+1} = \frac{(1 + A\Delta t \epsilon^2 k^4)\tilde{\phi}_i^{\,t} + \Delta t ik \cdot \Big( M(c^t) \Big[ ik' \Big( \frac{\partial \tilde{F}^t_{FH}}{\partial \phi_i} + \epsilon^2 k^2 \tilde{\phi}_i^{\,t} \Big) \Big]_r \Big)_k}{1 + A\Delta t \epsilon^2 k^4} \qquad [11]$$

we finally obtain our desired solution by taking the inverse Fourier transform and adding the noise term:

$$\phi_i^{t+1} = (\tilde{\phi}_i^{\,t+1})_r + \sigma \cdot \zeta \qquad [12]$$

Numerical implementation of the various spatial derivatives was guided by (16). We used an implicit time scheme to discretize the gelation equation:

$$\frac{c^{t+1} - c^t}{\Delta t} = M_c g(\phi_p^{t+1}, \phi_r^{t+1}) c^{t+1} - M_c c^t c^{t+1} \qquad [13]$$

Solving for $c^{t+1}$:

$$c^{t+1} = \frac{c^t}{1 - \Delta t M_c g(\phi_p^{t+1}, \phi_r^{t+1}) + \Delta t M_c c^t} \qquad [14]$$

Due to numerical difficulties associated with large values of the polymerization parameters $N_p$ and $N_r$ in the Flory-Huggins free energy, we used an approximation that was numerically tractable. Large values of $N_i$ result in numerically unresolvable small values for the dilute phase concentrations, so we smoothly interpolated the free energy near 0 with the case when $N_i = 1$ using the following function, $\hat{F}_{FH}$:

$$\begin{aligned} \hat{F}_{FH} = \Big[ \frac{\phi_p}{N_p}\Big(1 - e^{-\alpha\phi_p}\Big) + \phi_p e^{-\alpha\phi_p} \Big] \ln\phi_p + \Big[ \frac{\phi_r}{N_r}\Big(1 - e^{-\alpha\phi_r}\Big) + \phi_r e^{-\alpha\phi_r} \Big] \ln\phi_r + \\ (1 - \phi_p - \phi_r)\ln(1 - \phi_p - \phi_r) + \chi_{pr}\phi_p\phi_r + \chi_{pb}\phi_p(1 - \phi_p - \phi_r) + \chi_{rb}\phi_r(1 - \phi_p - \phi_r) \end{aligned} \qquad [15]$$

All simulations were run with $\Delta t = 0.0005$ for 4e6 timesteps, $\epsilon = 2$, $N_p = N_r = 30$, $\alpha = 30$, and $\sigma = 0.0001$. The gelation parameters were $c(t = 0) = \phi_p/1000$, $c_{\phi_p} = c_{\phi_r} = 0.01$, and $M_c = 0.2$.

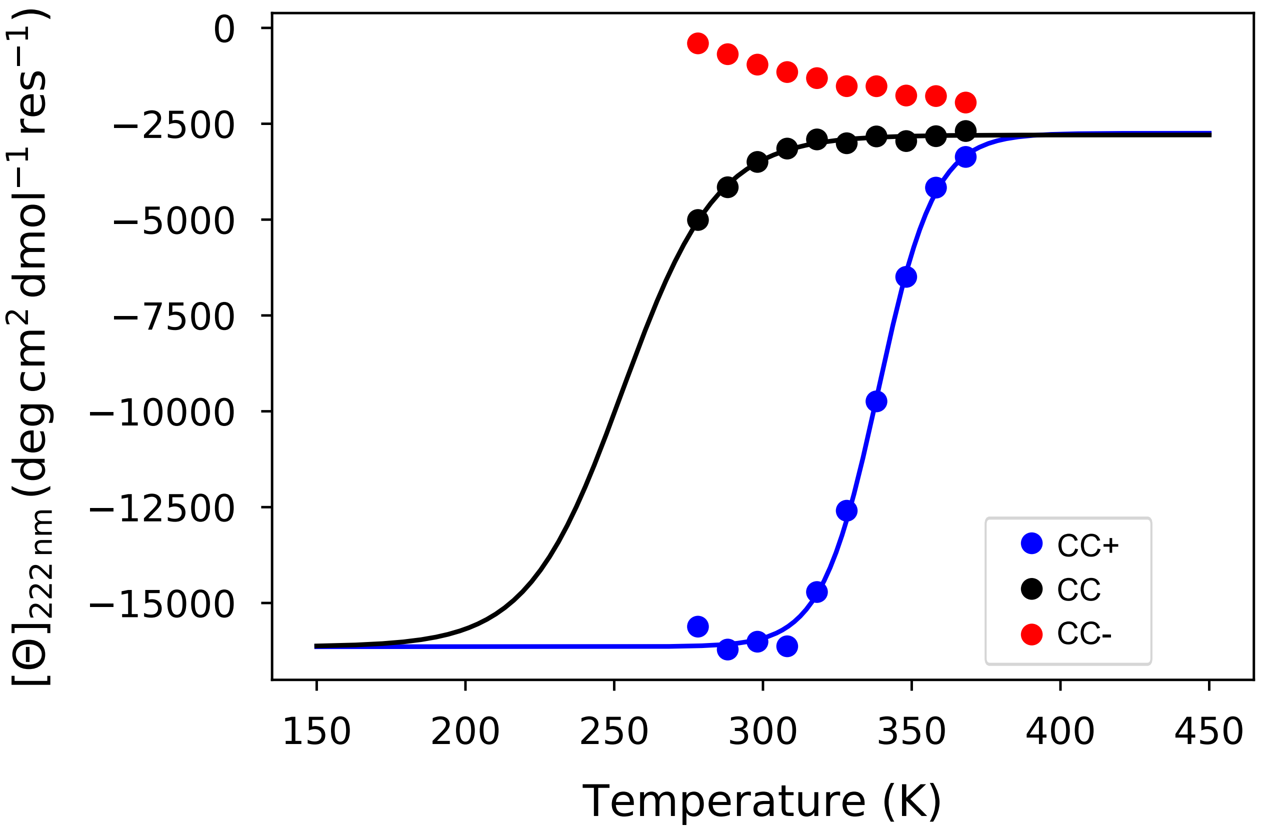


**Fig. S1.** Two-state model fits to the mean residue ellipticity at 222nm for each of the three peptides.

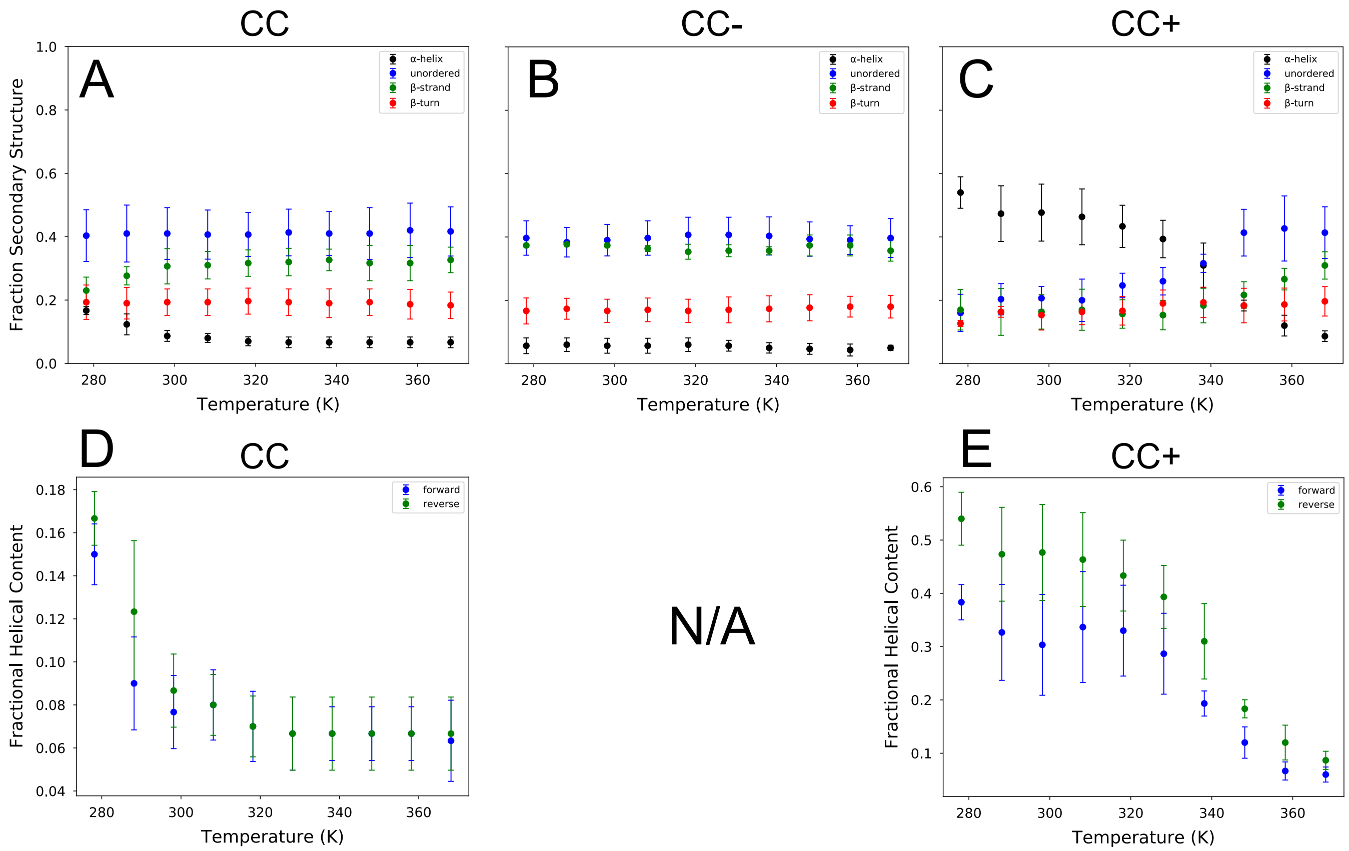


**Fig. S2. Secondary structure contents of each peptide for temperature-dependent CD** In all plots, secondary structure was determined by deconvolution using the CDSSTR algorithm and basis sets 4, 7, and SP175, as implemented by the Dichroweb server. Filled circles and error bars represent the mean and standard deviation, respectively, of the analysis resulting from these three basis sets. See SI Methods for details. (A) Secondary structure content for the WT peptide. (B) Secondary structure content for the CC- peptide. (C) Secondary structure content for the CC+ peptide. (D) Fractional helical content for the WT peptide for forward and reverse temperature CD scans. (E) Fractional helical content for the CC+ peptide for forward and reverse temperature CD scans.

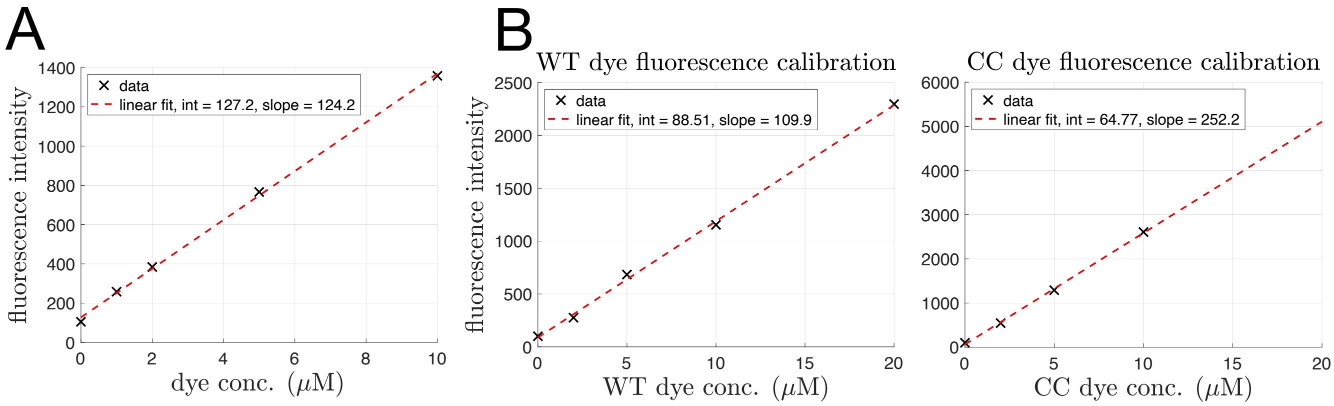


**Fig. S3.** Dye fluorescence calibration curves for measuring dense phase Whi3 concentrations. Fluorescence intensities of Whi3 in experiments fall within the ranges covered by the calibration curves.

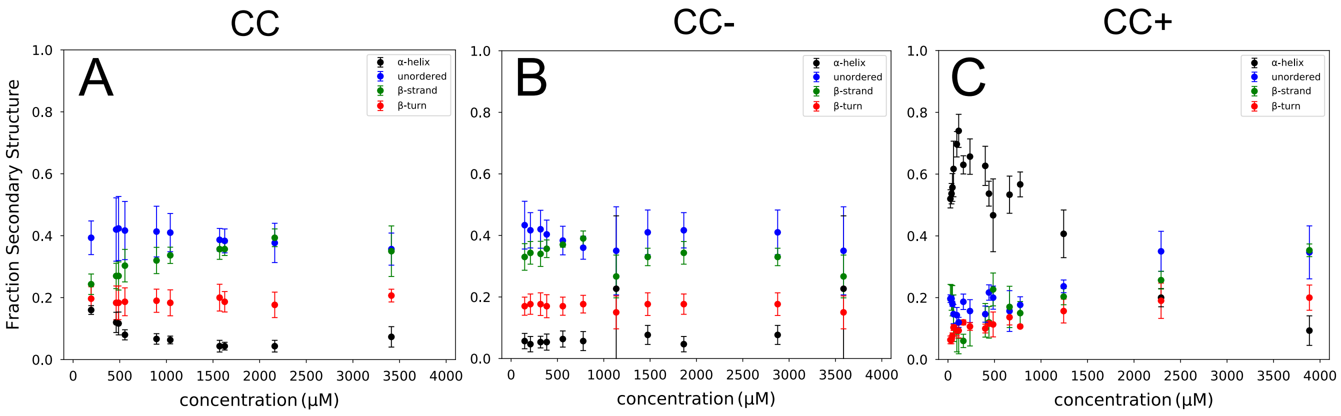


**Fig. S4. Secondary structure contents of each peptide for concentration-dependent CD** In all plots, secondary structure was determined by deconvolution using the CDSSTR algorithm and basis sets 4, 7, and SP175, as implemented by the Dichroweb server. Filled circles and error bars represent the mean and standard deviation, respectively, of the analysis resulting from these three basis sets. See SI Methods for details. (A) Secondary structure content for the WT peptide. (B) Secondary structure content for the CC- peptide. (C) Secondary structure content for the CC+ peptide.

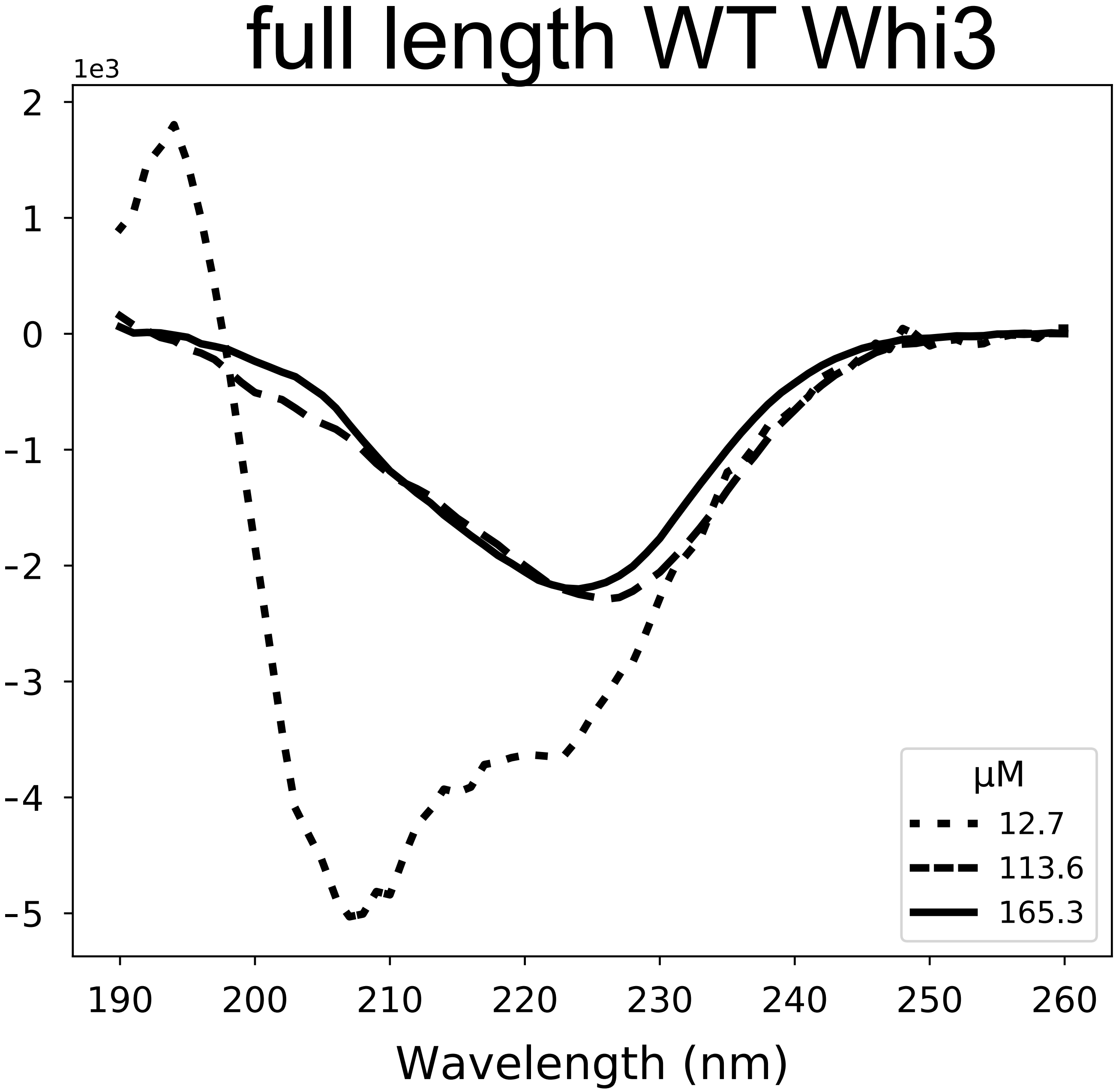


**Fig. S5.** CD spectra are shown for full-length WT Whi3 at three concentrations.

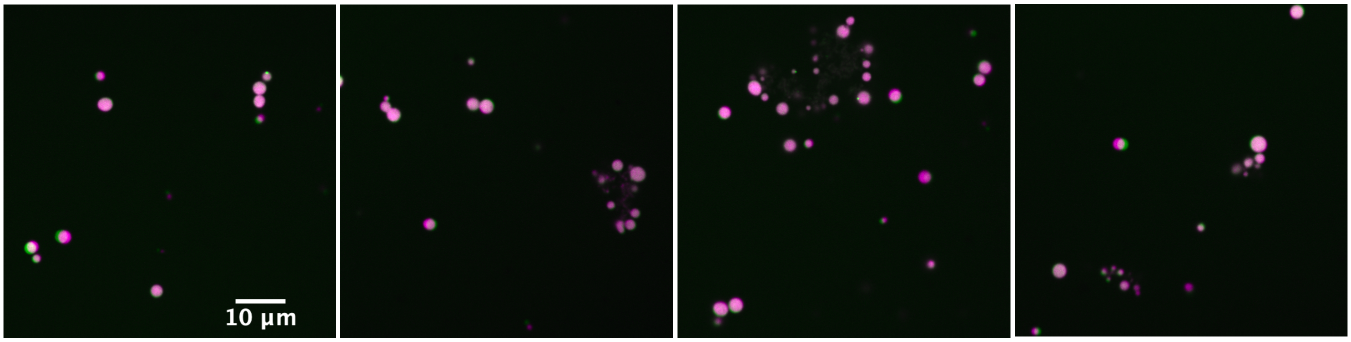


**Fig. S6.** Whi3 CC+ structures five hours after mixing 1$\mu$M Whi3 with 5nm *CLN3*. Images are contrasted identically to maximize visualization and do not reflect protein or RNA concentrations as in the main text.

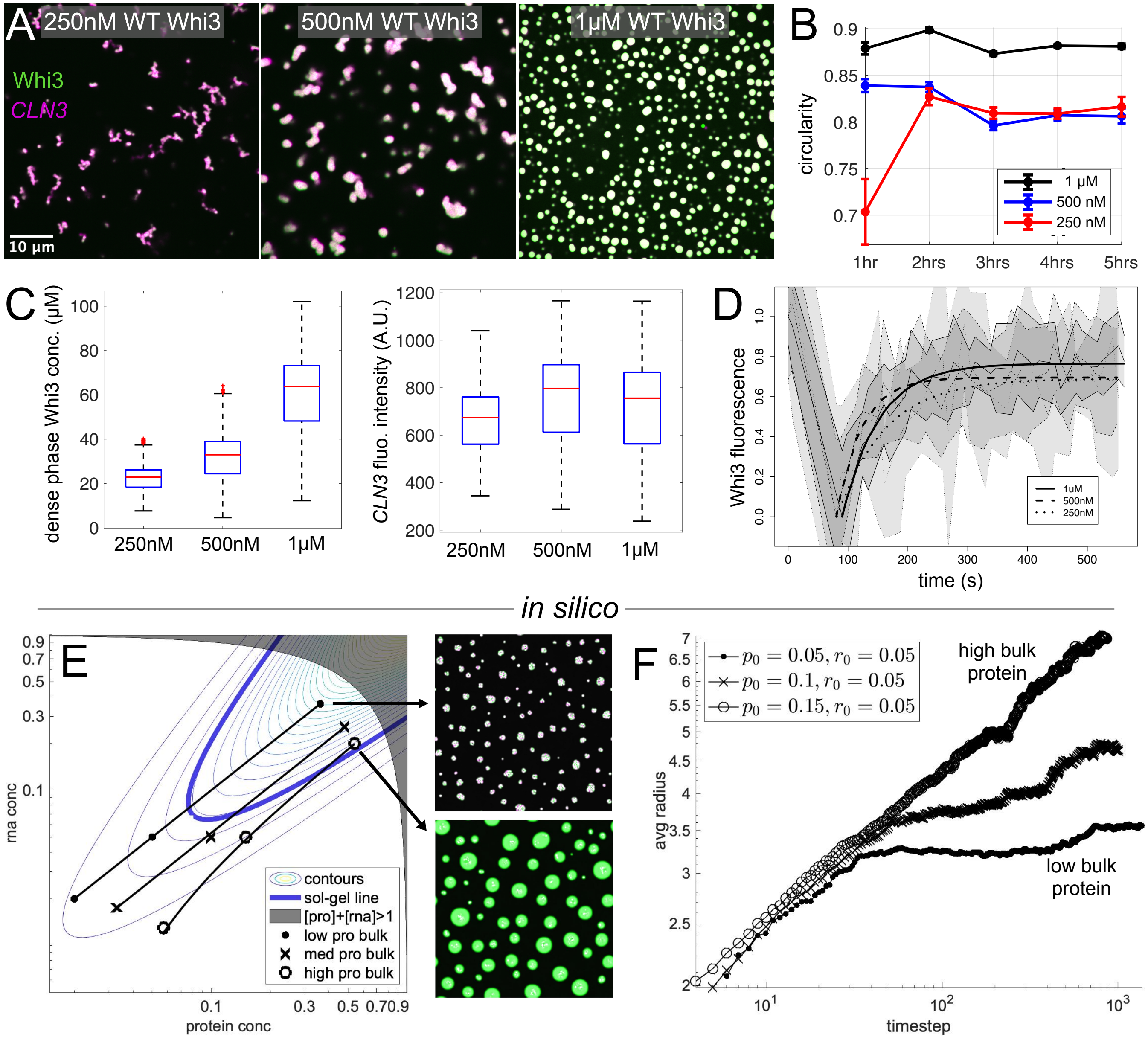


**Fig. S7. Bulk Whi3 concentration controls dense phase stoichiometry and dynamics.** (A) Representative images of WT Whi3 at different bulk concentration five hours after mixing with 5nM *CLN3*. Whi3 (green) and *CLN3* (magenta) channels are merged and contrasted identically. (B) Quantification of average droplet circularity for 3 replicates of experiments corresponding to panel (A). Error bars are standard error of the mean. (C) Dense phase Whi3 concentration and *CLN3* fluorescence intensity calculated for each condensate in experiments corresponding to panel (A). (D) Fluorescence recovery after photobleaching (FRAP) curves for WT Whi3 with *CLN3* at different bulk concentrations of Whi3 at around 2.5 hours after mixing. Solid lines show averages over 5-8 droplets each, with shaded regions indicating +- 1 standard deviation. (E) Parabolic sol-gel contours with the sol-gel line in bold blue are overlaid by dilute, bulk, and dense phase equilibrium concentrations of simulated systems initialized at different bulk protein concentrations but fixed interaction energies. Altered fixed bulk concentrations of protein result in dense phase concentrations that lie at varying depths beyond the sol-gel line. Simulation snapshots after steady state has been reach in the low bulk protein and high bulk protein concentration systems are shown. (F) The average radii of droplets from simulations in panel (E) are shown as a function of time. The legend provides values for the bulk concentrations of protein and RNA that determine whether the systems encode low, medium, or high bulk protein concentrations.

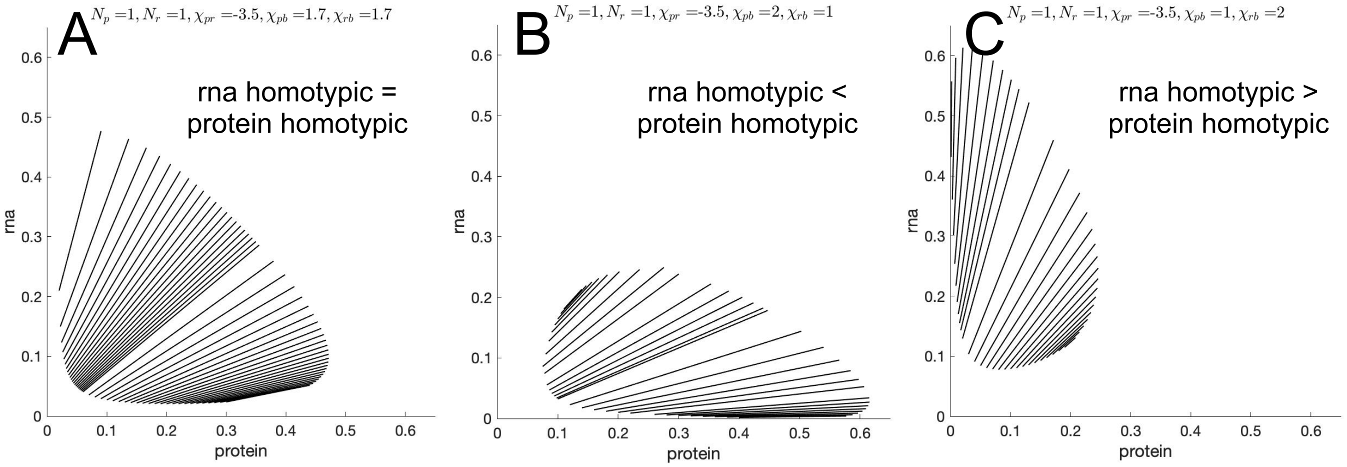


**Fig. S8. The balance of homotypic interactions determines the slopes of tie lines in FH ternary phase diagrams.** (A) When protein and rna homotypic interactions are equivalent, a phase diagram that is symmetric about the [rna]=[pro] line results. As more protein or rna is introduced to the system, the slopes of tie lines more closely match the respective axes. (B) When protein homotypic interactions dominate, the phase diagram is altered, and tie lines become more aligned with the protein axis. (C) When rna homotypic interactions dominate, the phase diagram is altered, and tie lines become more aligned with the rna axis. Note that since protein and rna homotypic interaction parameters are switched relative to panel (D), the phase diagram is simply reflected over the line [rna]=[pro].

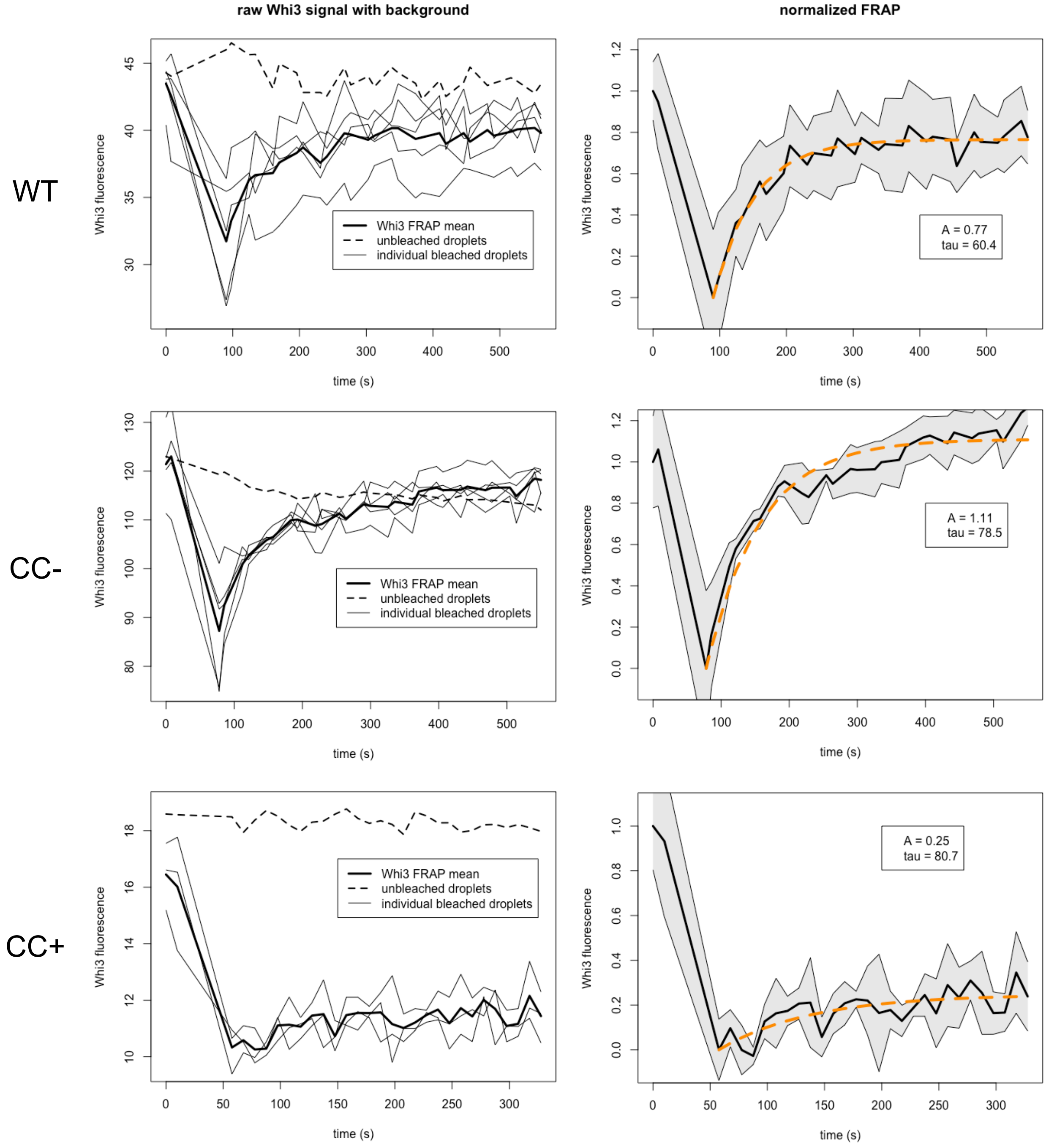


**Fig. S9. FRAP experiment raw data, normalizations, and model fits.** In the left column, for each experiment, the raw Whi3 fluorescence signal for each bleached droplet is shown, along with their mean and the mean fluorescence signal of the unbleached droplets in the same field of view. In the right column, the normalized signal is shown along with the model fit and fit parameters.

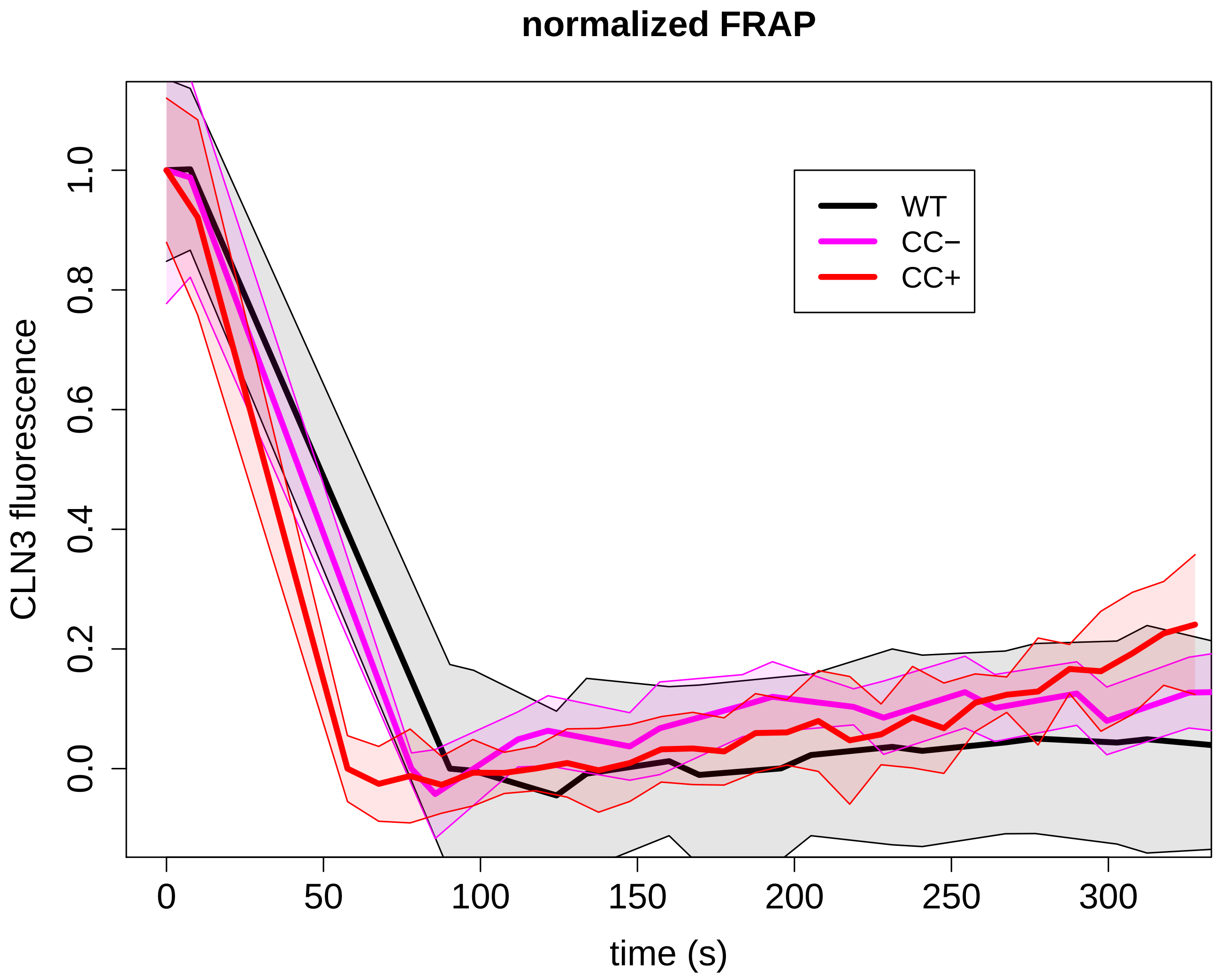


**Fig. S10.** Normalized fluorescence recovery after photobleaching (FRAP) curves plus and minus 1 standard deviation for *CLN3* mixed with Whi3 CC variants 2 hours after mixing.

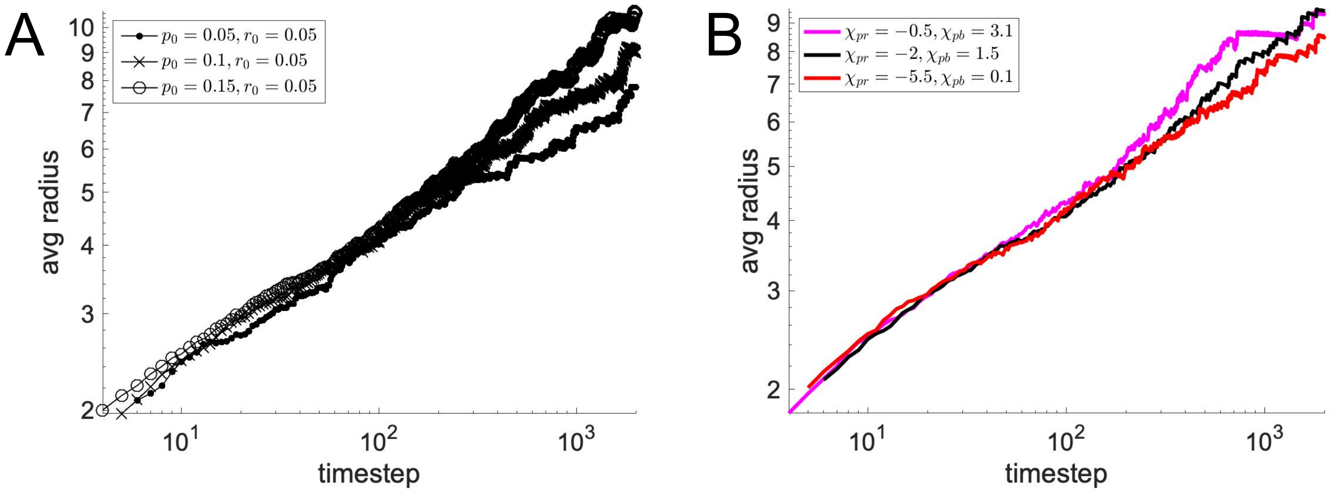


**Fig. S11. Average droplet size is not significantly affected without gelation.** (A) Average droplet size as bulk concentration of protein is increased. At long times when each system is a single droplet, the average droplet radius will differ. (B) Average droplet size as bulk concentrations are fixed, but interaction parameters change.

**Table S1. Whi3 and *CLN3* constructs**

| Primer Name | Construct | Sequence |
|---|---|---|
| AGO3201 | WT, CC-, CC+, ΔCC | ctttaagaaggagatatacatatgcaccatcatcatcatcatatgtcgctggttaacagt |
| AGO3024 | WT, CC-, CC+, ΔCC | aagcttgtcgacggagctcgtcaagatttgccgaaggc |
| AGO3086 | ΔCC | ATGCTGGTGATGCTGGAGTGCTTGggattgtggc |
| AGO3090 | ΔCC | gccacaatccCAAGCACTCCAGCATCACCAGCATcagcagcaacagt ctctttc |
| AGO3085 | CC- | gGGTGGTGCTGCTGCTGCTGGgGTTGCTGCTGCTGGTG ATGCggatgctggtgatgctgg |
| AGO3084 | CC- | ccGCATCACCAGCAGCAGCAACcCCAGCAGCAGCAGCAC CACCc-CCAGCAGCAGCAGCAC |
| AGO3087 | CC+ | tGCAGCAGCAACTCCAGCAGCtGCAGCACCACCTCCAG CAGCtGCAGCACCAGCAG |
| AGO3088 | CC+ | aGCTGCTGGAGGTGGTGCTGCaGCTGCTGGAGTTGCTG CTGCaGGTGATGCAAat-gctgg |
| AGO3001 | CLN3 | TAATACGACTCACTATAGGGAGAGTCTGCATACCAAAGA TCAGCCGCTTGC |
| AGO3002 | CLN3 | GTATGCTAGCGTAGTTTCTTGACC |

**Table S2. LASSI simulation energies**

| **WT isotropic** | CC | QRR |
|---|---|---|
| CC | -0.1 | -0.1 |
| QRR | -0.1 | -0.2 |

| **WT anisotropic** | CC | QRR |
|---|---|---|
| CC | -2 | 0 |
| QRR | 0 | 0 |

| **CC+ isotropic** | CC | QRR |
|---|---|---|
| CC | 0 | 0 |
| QRR | 0 | -0.2 |

| **CC+ anisotropic** | CC | QRR |
|---|---|---|
| CC | -4 | 0 |
| QRR | 0 | 0 |

| **CC- isotropic** | CC | QRR |
|---|---|---|
| CC | -0.2 | -0.2 |
| QRR | -0.2 | -0.2 |

| **CC- anisotropic** | CC | QRR |
|---|---|---|
| CC | 0 | 0 |
| QRR | 0 | 0 |
